# An RNA-binding protein secreted by *Listeria monocytogenes* activates RIG-I signaling

**DOI:** 10.1101/543652

**Authors:** Alessandro Pagliuso, To Nam Tham, Eric Allemand, Stevens Robertin, Bruno Dupuy, Quentin Bertrand, Christophe Bécavin, Mikael Koutero, Valérie Najburg, Marie-Anne Nahori, Fabrizia Stavru, Andréa Dessen, Christian Muchard, Alice Lebreton, Anastassia V. Komarova, Pascale Cossart

**Author notes:** These authors contributed equally to this work (To Nam Tham, Eric Allemand, Stevens Robertin).

## Abstract

RNA binding proteins (RBPs) perform key cellular activities by controlling the function of bound RNAs. The widely held assumption that RBPs are strictly intracellular has been challenged by the discovery of secreted RBPs. While extracellular RBPs have been described in mammals, secreted bacterial RBPs have not been reported. Here, we show that the bacterial pathogen *L. monocytogenes* secretes a small RBP that we named Zea. We show that Zea binds a subset of *L. monocytogenes* RNAs causing their accumulation in the extracellular medium. Furthermore, during *L. monocytogenes* infection, Zea binds RIG-I, the non-self-RNA innate immunity sensor, potentiating interferon β production. Mouse infection studies revealed that Zea modulates *L. monocytogenes* virulence. Together, this study uncovered the presence of an extracellular ribonucleoprotein complex from bacteria and its involvement in host-pathogen crosstalk

---

RNA binding proteins (RBPs) are found in all living organisms. By binding RNAs, RBPs assemble in ribonucleoprotein complexes that dictate the fate and the function of virtually every cellular RNA molecule. In bacteria, RBPs interact with their cognate RNAs *via* classical RNA binding domains (RBDs), structurally well-defined signatures that recognize specific RNA sequences and/or motifs [reviewed in (*1*)]. Previously thought to be mainly involved in transcriptional regulation, bacterial RBPs have now been implicated in a wide variety of cellular processes such as translation, RNA turnover and decay, as well as RNA processing and stabilization (*1*). Although bacterial RBPs regulate vital functions in bacterial physiology, their number remains limited. This is mainly due to the difficulty to apply in bacteria the recently developed proteomic approaches that greatly expanded the number of RBPs in eukaryotes (*2, 3*). These approaches almost doubled the eukaryotic census of RBPs and revealed the presence of hundreds of RBPs lacking classical RBDs. It is therefore conceivable that many bacterial RBPs remain to be discovered.

One feature of all so far described bacterial RBPs is their exquisite intracellular localization. Few extracellular RBPs have only been described in mammals and have been shown to stabilize RNA in the extracellular *milieu* and participate in cell-to-cell communication (*4–7*). At present, no secreted RBPs have been identified in bacteria. A recent study screened thousands of secreted effectors of Gram-negative symbionts and bacterial pathogens for the presence of known RBDs and failed to unambiguously identify any RBPs (*8*). It is therefore likely that secreted bacterial RBPs harbor unconventional RBDs which render them undetectable by using conservation-based searches.

In this study, we report the identification of the first secreted bacterial RBP, the *Listeria monocytogenes (L. monocytogenes)* protein Lmo2686. We provide evidence that Lmo2686 is secreted in the culture supernatant where it is associated with a subset of *L. monocytogenes* RNAs. Protein sequence analysis of Lmo2686 revealed the absence of a canonical RBD, suggesting a novel mode of RNA binding. We show that Lmo2686 induces the extracellular accumulation of its RNA targets possibly by protecting them from degradation. Furthermore, during infection of mammalian cells, Lmo2686 interacts with RIG-I and potentiates RIG-I-dependent type I interferon (IFN) response. We further show that Lmo2686 regulates *L. monocytogenes* virulence *in vivo*. Based on these findings, we propose to rename this protein Zea, as Zea, also known as Hecate, is an ancient Greek goddess who protected and guided the travelers. The presence of Zea orthologs in other bacterial species revealed that secretion of RBPs is a conserved phenomenon in prokaryotes.

## Zea is a small secreted protein of *L. monocytogenes*

*The lmo2686/zea* open reading frame is 534-bp long and is located 164 bp downstream from the *lmo2687* gene, which encodes an FtsW-related protein. *zea* is followed by a transcriptional terminator upstream from the divergently transcribed *lmo2685* gene, which encodes a cellobiose phosphotransferase protein (**Figure 1A**). *zea* is found in about half of the *L. monocytogenes* strains sequenced to date as well as in the animal pathogen *Listeria ivanovii* (*9*). Orthologs of *zea* are also found in other species, mainly Gram-positive bacteria of the genus *Bacillus* (**Figure S1**). Interestingly, *zea* is absent from the genome of the nonpathogenic species *L. innocua* (*10*) and *Listeria marthii* FSL S4-120 (*11*) which suggests that it may play a role in *L. monocytogenes* virulence (**Figure 1A**).

**Figure 1.**
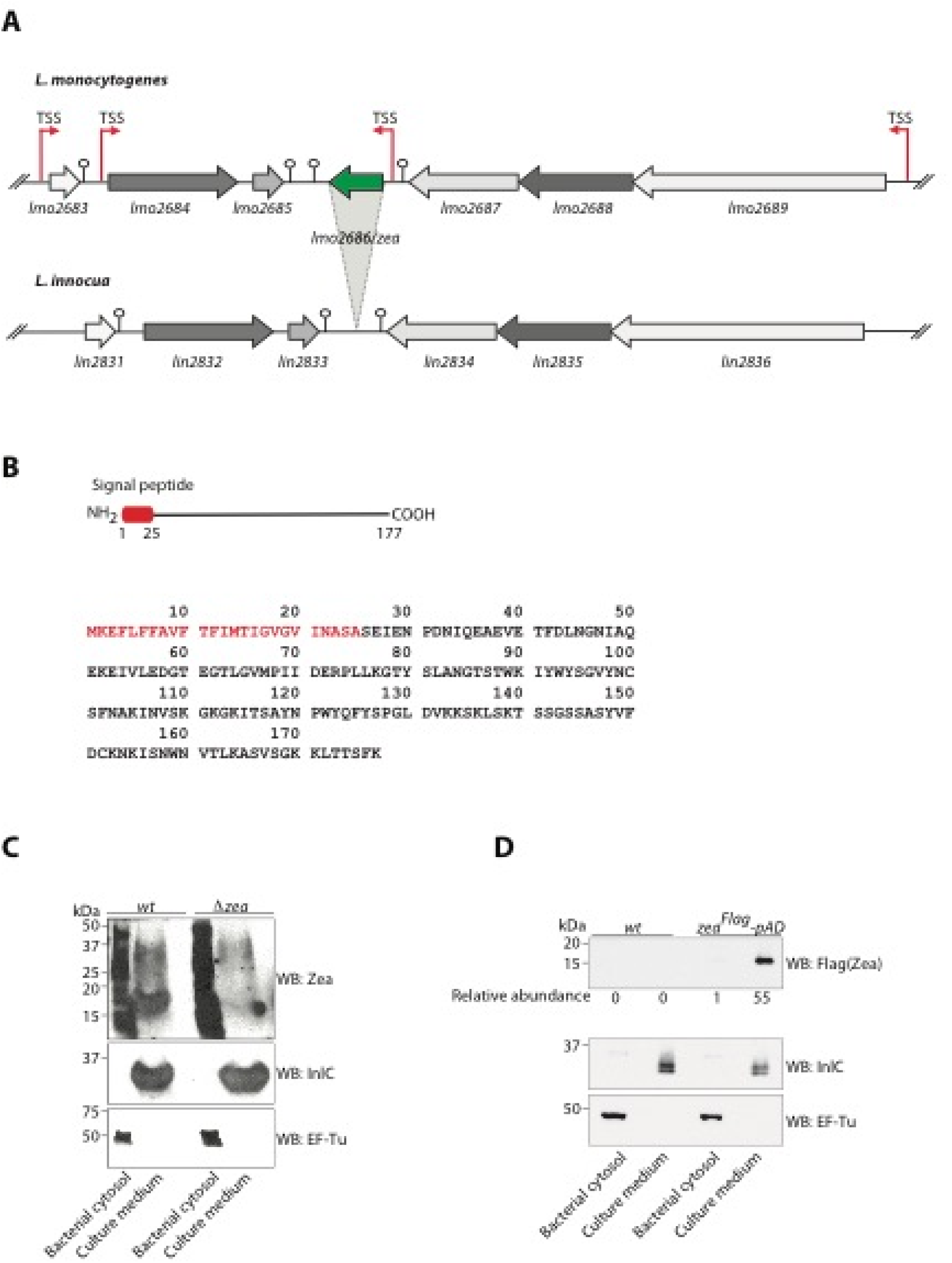
Zea is a secreted *L. monocytogenes* protein. (**A**) Syntheny analysis of the *lmo2686/zea*-containing genomic locus between *L. monocytogenes* and *L. innocua* showing the absence of the *lmo2686/zea* gene in *L. innocua.* Arrows represent the transcriptional start sites (TSS). The stem and circle represent the transcriptional terminators. (**B**) Schematic representation (top) and primary sequence (bottom) of the Zea protein. The N-terminal signal peptide of 25 amino acids is highlighted in red. (**C**) Bacterial cytosol and culture medium were prepared from *L. monocytogenes wt* and *Δzea* strains and immunoblotted with the indicated antibodies. EF-Tu and InlC were used as markers of the bacterial cytosol and culture medium, respectively. (D) Bacterial cytosol and culture medium from *L. monocytogenes wt* and a Flag-tagged Zea-overexpressing *L. monocytogenes* strain *(zea^Flag^-pAD)* were immunoblotted with the indicated antibodies. Values of the relative abundance of ZeaFlag for one representative western blot were generated with ImageJ.

RNA sequencing (RNA-Seq) data has revealed the presence of a transcriptional start site upstream of the start codon of *zea* which indicates that the gene is transcribed (**Figure 1A**) (*12*). *zea* appears constitutively expressed at 37 °C, albeit at low levels, and is slightly more expressed at 30 °C and under microaerophilic conditions (*9, 12*).

The *zea* gene encodes a small protein of 177 amino acids (aa) (**Figure 1B**). Analysis of the Zea protein sequence predicted the presence of a N-terminal signal peptide of 25 aa for SecA-mediated secretion, resulting in a putative 152 aa mature protein with a basic isoelectric point (pI = 8.4) (**Figure 1B**). We could not identify any other domain or sequence of known function.

The presence of a signal peptide prompted us to test whether Zea could be secreted. We generated three different antibodies against three peptides of the C-terminus of the protein and used them to assess the presence of Zea in the *L. monocytogenes* cytosol and in the culture medium. Western blotting analysis revealed that Zea could be recovered from the culture medium, indicating secretion of the protein (**Figure 1C**). Culture medium collected from the *zea-deleted strain (Δzea)* did not show any immunoreactive band, thus confirming the specificity of our antibodies. The secretion of Zea was also confirmed by engineering a *L. monocytogenes* strain carrying a chromosome-integrated copy of the C-terminally Flag-tagged *zea* gene under the control of a constitutive promoter (*zea^Flag^-pAD* (**Figure 1D**). Quantitative analysis of the distribution of Zea between bacterial cytosol and culture medium in stationary phase revealed a strong accumulation in the extracellular medium indicating that the protein was efficiently secreted (**Figure 1D**).

## Zea is an oligomeric protein that interacts with RNA

The structure of Zea was previously solved by X-ray crystallography at a resolution of 2.75 Å and deposited in the Protein Data Bank (PDB) by Minasov G. and colleagues (PDB ID: 4K15; https://www.ebi.ac.uk/pdbe/entry/pdb/4K15). According to this structure, Zea is a toroid-shaped homohexamer in which every monomer contacts the neighboring molecule *via* a beta-hairpin-beta unit (**Figure 2A**). As this structure is not shared by any other polypeptide of known function, at present, the role of Zea and of its orthologs in other species is unknown.

**Figure 2.**
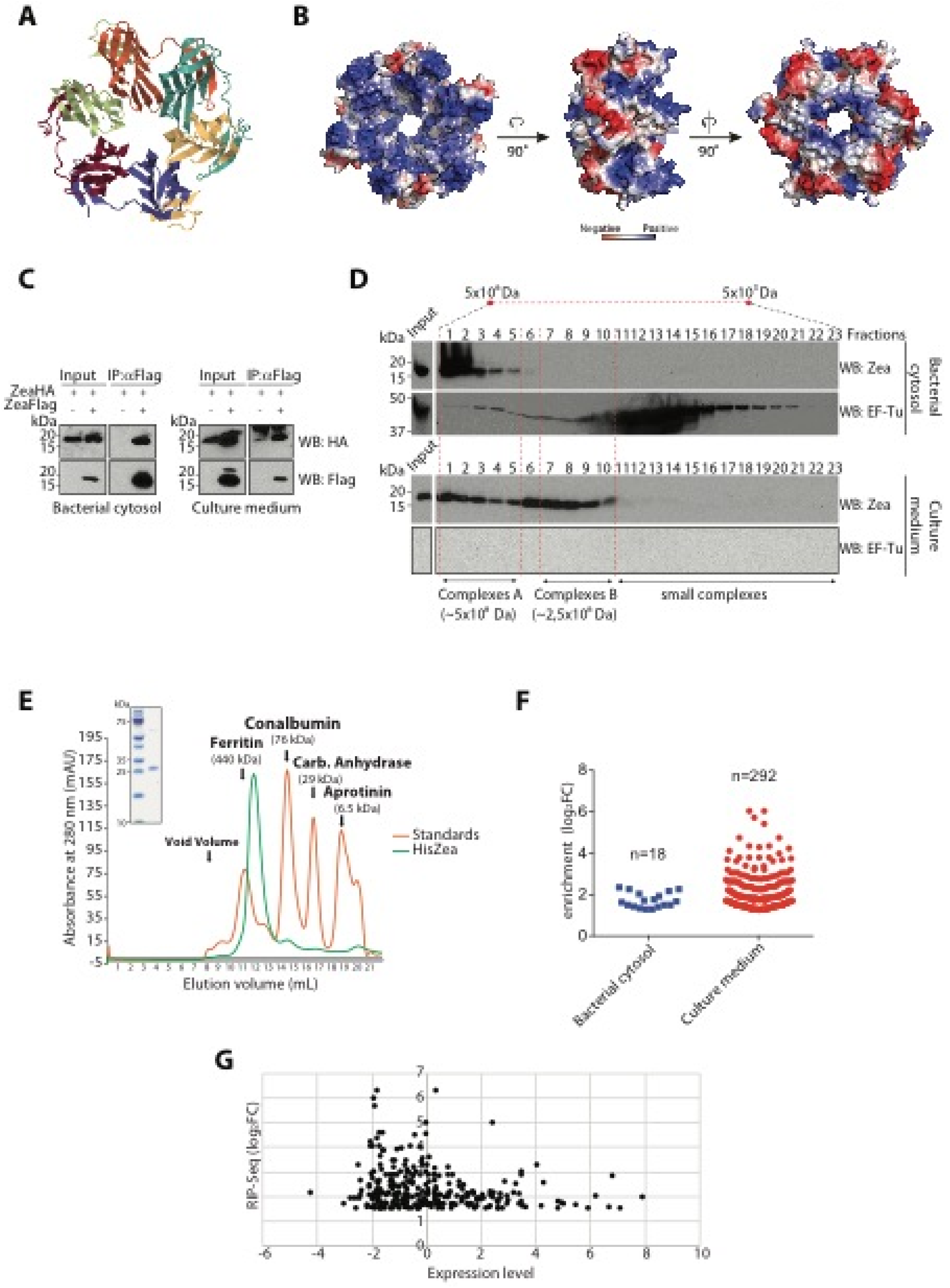
Zea is an oligomeric protein that interacts with a subset of *L. monocytogenes* RNAs. **(A)** Ribbon diagram of hexameric Zea. Every monomer is depicted with a different color. **(B)** Electrostatic potential surface representation of the Zea hexamer. The molecule is rotated by 90 °C about the y-axis in the successive images. **(C)** Immunoprecipitation (IP) of Zea with an anti-Flag antibody from bacterial cytosol and culture medium from a *L. monocytogenes* strain co-overexpressing ZeaFlag and ZeaHA. The starting material (Input) and Flag-immunoprecipitated proteins (IP: αFlag) were probed by western blotting (WB) with an anti-Flag and with an anti-HA antibodies: starting material (Input) and Flag-immunoprecipitated proteins (IP: αFlag). **(D)** ZeaFlag elution from size exclusion gel chromatography. *L. monocytogenes* bacterial cytosol and culture medium were applied to Superose 6 size exclusion gel chromatography column. An aliquot of each fraction was precipitated with acetone and analyzed by WB with the indicated antibodies. The cytosolic protein EF-Tu serves as a control for the absence of intact or lysed bacteria in the culture medium fraction. **(E)** 280 nm (mAU) absorbance monitoring of a gel filtration profile (Superdex 200) of recombinant purified HisZea (green line). The elution profile of protein markers of known molecular weight is indicated with the orange line. HisZea was purified by Ni^2+^ -affinity followed by size exclusion gel chromatography, analyzed by SDS-PAGE and Coomassie blue staining (top left-hand panel). **(F)** Enrichment of Zea-bound RNAs (n) from bacterial cytosol and culture medium. Blue squares and red circles depict individual RNAs. The y-axis shows the enrichment (expressed as log_2_FC) of the Zea-interacting RNAs relative to immunoprecipitation with preimmune serum. **(G)** Expression of *L. monocytogenes* RNAs grown in BHI at stationary phase measured by tiling array compared to the enrichment of the Zea-bound RNAs (expressed as log2FC).

We noticed that several other proteins that assemble as a torus, like Zea, have the intrinsic capability to bind RNA (*13–17*). Interestingly, Zea shows a positively charged surface on one side of the torus, due to the presence of several lysine residues, which might well accommodate the negatively-charged RNA (**Figure 2B**). These features led us to hypothesize that Zea might bind RNA.

Before addressing this hypothesis, we sought to verify whether the oligomeric state of Zea observed by X-ray crystallography also existed under physiological conditions, ruling out possible crystallization artifacts. We used three different approaches: (i) co-immunoprecipitation of HA- and Flag-tagged versions of Zea (**Figure 2C**), (ii) size exclusion chromatography of *L. monocytogenes* cytosol and culture medium (**Figure 2D**) and (iii) size exclusion gel chromatography of recombinant His-tagged Zea expressed and purified from *E. coli* (**Figure 2E**). Collectively, our data show that Zea has a high tendency to oligomerize, in line with the hexameric structure shown by X-ray crystallography.

We then examined whether Zea could bind RNA. We performed RNA immunoprecipitation (IP) of cytosolic extract and culture supernatant followed by high-throughput sequencing (RIP-Seq) by using the protocol schematized in **Figure S2**. Given the low amount of Zea protein produced *in vitro,* we made use of a Zea-overexpressing strain *(zea-pAD).* Zea was immunoprecipitated from *L. monocytogenes* cytosol and culture medium and the Zea-bound RNAs were subsequently extracted and sequenced. As a control, we performed a mock IP, using an unrelated antibody of the same isotype. Remarkably, RIP-Seq analysis revealed the presence of several *L. monocytogenes* RNAs, which were highly enriched compared to the control samples (**Figure 2F**), indicating that Zea can form complexes with RNA. An enrichment threshold of log2 fold change (log_2_FC) >1.5 (corresponding to an almost three-fold increase) was used for the identification of Zea-associated RNAs. This filtering led to the identification of 18 and 292 RNAs bound to intracellular and secreted Zea, respectively (**Figure 2F**). The enrichment of Zea-bound RNAs was particularly high in the culture medium fraction with almost 30% of the identified RNAs showing a log_2_FC ranging from 2.5 up to 6 (five to sixty times more present in the Zea IP compared to the control IP). Importantly, the enrichment of specific RNAs in the Zea IP was uncorrelated to their expression levels (**Figure 2G**) (*9*). Together, our data suggest that Zea binds specific RNA targets. Indeed, we could not detect binding of the most highly expressed – and most stable – RNAs, such as rRNAs and tRNAs. In contrast, we found that Zea bound preferentially a subset of protein-coding mRNAs and to lesser extent small regulatory RNAs (**Table S1**).

We then analyzed the genomic distribution of Zea-bound RNAs on the *L. monocytogenes* chromosome (**Figure 3A**). It was striking that there was one region particularly over-represented. This locus contains the prophage A118 (**Figure 3A-C**), and the phage RNA was present both in the bacterial cytosol and in the culture medium (**Table S1**). Cluster of Orthologous Genes (COG) classification highlighted the phage A118 RNA as the most enriched class of Zea-bound RNAs (**Figure S3**). The phage A118 is a temperate phage belonging to the *Syphoviridae* family of double-stranded DNA bacterial viruses (*18*).

**Figure 3.**
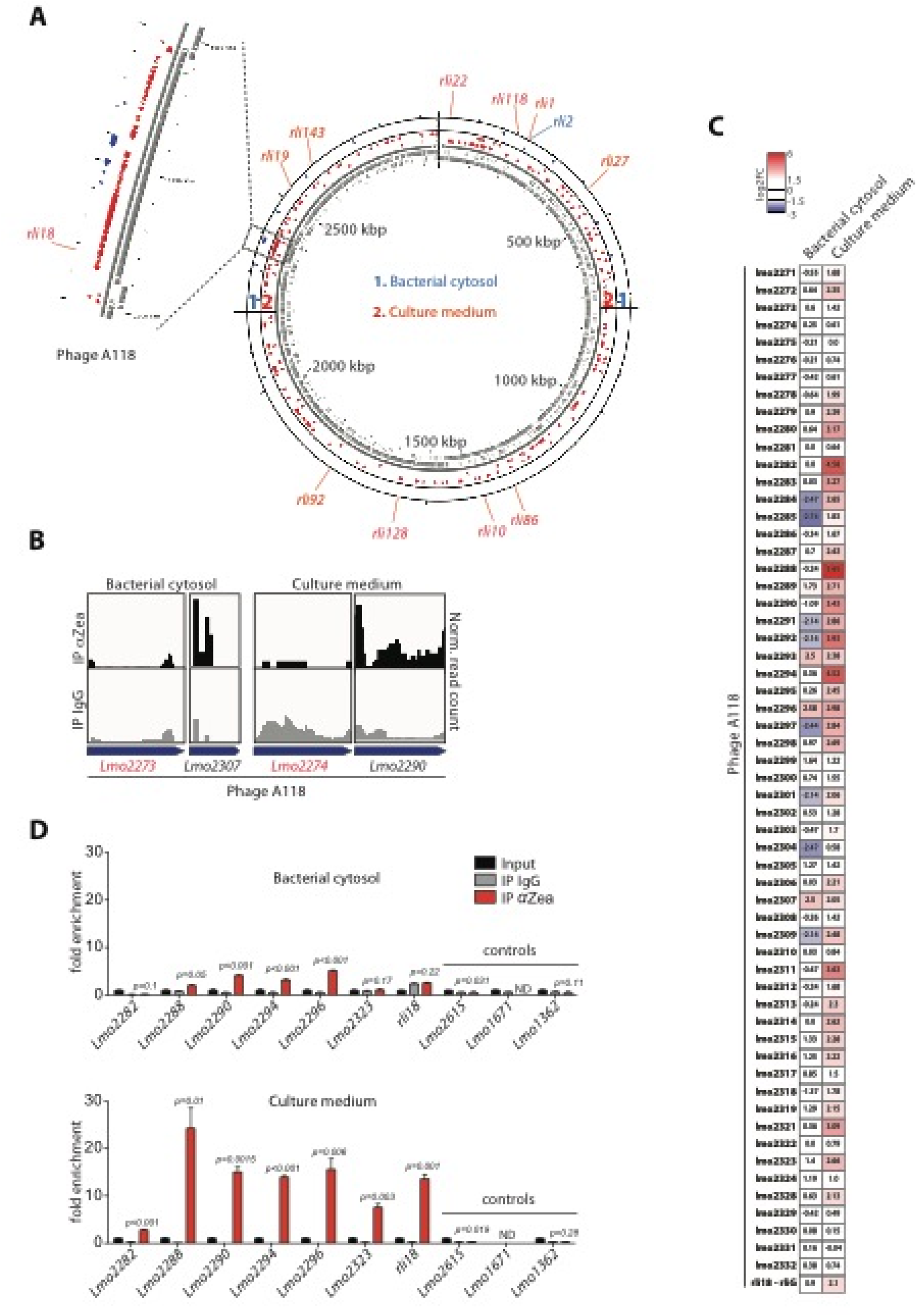
Zea interacts with phage RNA. **(A)** Circular genome map of *L. monocytogenes* showing the position of the Zea-interacting RNAs. The first two circles from the inside show the genes encoded on the + (inner track) and – (outer track) strands, respectively. The positions of Zea-interacting small RNAs (rlis) are pointed at outside of the circular map. Dotted lines highlight the positions of the phage A118 in the *L. monocytogenes* genome. **(B)** Examples of normalized read coverage (reads per million), visualized by IGV from Zea and control (IgG) immunoprecipitations (IP) for a selection of phage A118 phage genes. Genes are depicted with blue arrows. Gene names marked in red show no significant enrichment in the Zea IP compared to IgG IP. **(C)** Heat map showing the fold enrichment of phage A118 transcripts in the Zea IP compared to control IP, in the bacterial cytosol and culture medium for the phage A118 (values are expressed as log2FC). **(D)** RIP-qPCR on RNAs isolated from Zea and control (IgG) immunoprecipitations in the bacterial cytosol (top panel) and culture medium (bottom panel). The enrichment of selected phage *(lmo2282* to *lmo2333*) and control genes was calculated after normalization to the corresponding input fractions. Values reprents means ± SEM, n=3. ND, not detected. Statistical significance (between the IP IgG and IP αZea) is determined by two tailed t-test.

To validate the interaction of phage A118 RNA with Zea found by RIP-Seq (**Figure 3A-C**), we next performed RNA IP coupled to quantitative PCR (RIP-qPCR) analysis. RIP-qPCR confirmed a strong association of Zea to phage RNA, while we could not find any binding to control transcripts that were not enriched in our RIP-Seq dataset (**Figure 3D**). In agreement with the RIP-Seq data, the enrichment of phage RNA in the culture medium fraction was particularly strong (up to approximately 30 times more compared to control IgG IP) indicating that the phage RNA accumulates extracellularly together with Zea (**Figure 3D**). Taken together, our data revealed that Zea is an oligomeric protein that binds a subset of *L. monocytogenes* RNAs both in the bacterial cytosol and in the culture medium.

## Zea directly binds *L. monocytogenes* RNA

We next investigated whether Zea could directly bind RNA. We first performed electrophoretic mobility gel shift assay (EMSA) by using recombinant His-tagged Zea (HisZea) and *in vitro-* transcribed radiolabeled RNA. We selected *rli143* and *rli92,* two small RNAs that showed a significant enrichment in the RIP-Seq dataset (eight- and almost three-fold enrichment compared to control IgG IP, respectively; see **Figure 3A**). Incubation of *rli143* or *rli92* with HisZea produced several obvious shifts, which are likely due to the binding of different Zea oligomers to RNA (**Figure 4A-B**). Importantly, the binding of Zea with both *rli143* and *rli92* was specific, as it was displaced by the addition of increasing amounts of each unlabeled small RNA (**Figure 4C-D**).

**Figure 4.**
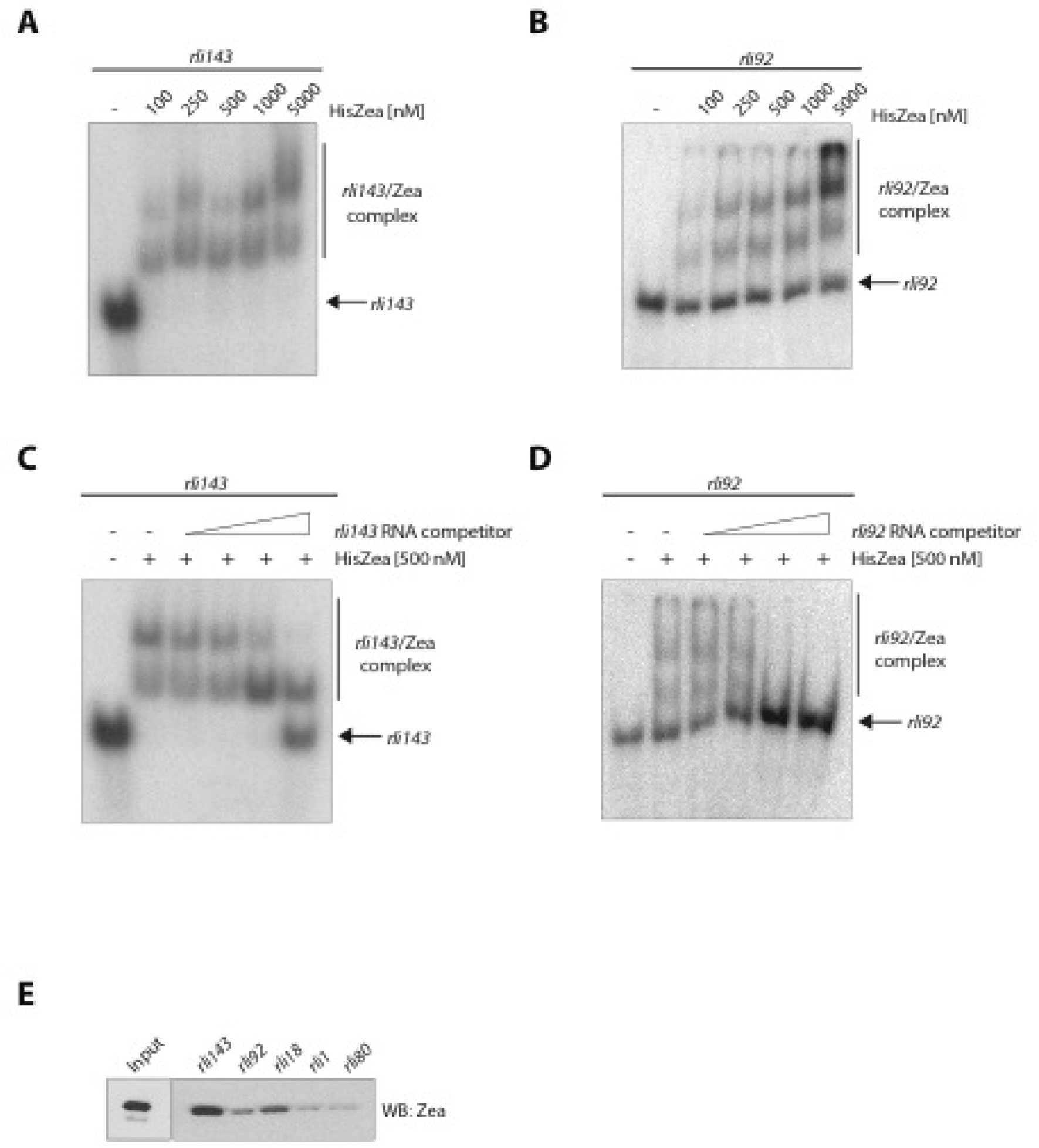
Zea directly binds RNA. **(A, B)** Representative electrophoretic mobility gel shift assay (EMSA) with *in* vitro-transcribed 5’-end radiolabeled *rli143* and *rli92* in the presence of increasing concentration of HisZea, as indicated. **(C, D)** HisZea-*rli143* and HisZea-*rli92* complexes were incubated with increasing concentrations of the corresponding cold competitor RNA. **(E)** Representative WB of streptavidin affinity pull-down of *in vitro-tran*scribed biotinylated transcripts (500nM) in the presence of HisZea (400nM). Zea input, 100 ng.

To further prove direct binding of Zea with its target RNAs, we performed an RNA pull-down assay, using *in* vitro-transcribed biotinylated RNA and recombinant HisZea. Here, in addition to *r1i143* and *r1i92*, we tested two other small RNAs (*r1i18* and *r1i1*), which also displayed specific binding to Zea in the RIP-Seq dataset (**Figure 3A**). As a control we employed a small RNA (rli’80), that was not specifically bound by Zea. Biotinylated small RNAs were incubated with HisZea and then recovered by streptavidin-coupled beads. After extensive washing, the amount of bound HisZea was assessed by western blotting. Using this approach, we detected binding above background level for three of the four small RNAs tested (**Figure 4E**). Importantly, *rli143,* which showed the highest enrichment in the RIP-Seq dataset, was also the strongest Zea binder in both the gel shift assay and the pull-down experiments, further validating the RIP-Seq results. Collectively, these data clearly indicate that Zea can stably and directly bind RNA.

## Extracellular Zea-bound RNAs do not derive from bacterial lysis

It was of importance to verify whether the extracellular RNAs in complex with Zea were not due to bacterial lysis. We thus first analyzed the presence of one of the most abundant cytosolic proteins of *L. monocytogenes* (EF-Tu) in the culture medium. While EF-Tu was readily detected in the bacterial cytosol, it was undetectable in the culture medium, indicating that bacterial lysis under our experimental conditions was negligible (**Figure 2D**). As bacteria lyse after death, we also quantified live and dead bacteria in our experimental conditions. Confocal microscopy analysis revealed very few dead bacteria (less than 2%), further confirming minimal bacterial lysis (**Figure S4A**). Finally, we designed an experiment in which we used the strict intracellular localization of the RBP Hfq and of its RNA targets as a readout of bacterial lysis. In *L. monocytogenes,* Hfq has been shown to bind three small RNAs, *LhrA, LhrB* and *LhrC* (*19*). We reasoned that, if bacterial lysis occurred, we should find Hfq complexed to its RNA targets in the medium. To assess this, *L. monocytogenes* was grown under the conditions used for the Zea RIP-Seq experiment and Hfq was then immunoprecipitated from the bacterial cytosol and culture medium by using a specific anti-Hfq antibody (*19*). Hfq was recovered from the bacterial cytosol but was undetectable in the culture medium, indicating minimal bacterial lysis (**Figure S4B**). RNA was extracted from the immunopurified Hfq ribonucleoprotein complexes and used to assess the abundance of the Hfq targets by qPCR. Given the low expression of *LhrB* and *LhrC* in stationary phase (*19*), we focused on *LhrA. LhrA* was detected in association with intracellular Hfq, but remained undetectable in the culture medium (**Figure S4C**). Collectively, these results strongly indicate that extracellular RNAs complexed with Zea are not originating from lysed bacteria.

## Zea overexpression induces extracellular accumulation of Zea-binding RNAs

Since Zea binds RNA and is secreted, we sought to determine whether it could affect the amount of RNA in the culture medium. Given the strong binding of Zea to phage RNA (**Figure 3A-D and S3**), we compared the amount of phage RNA present in the culture medium of *L. monocytogenes wt, Δzea* and *zea-pAD.* For this purpose, *L. monocytogenes* was grown in minimal medium (MM), because rich medium (BHI) contains large quantities of contaminating RNA. qPCR analysis on RNA extracted from the culture medium revealed that the overexpression of Zea increased the amount of the extracellularly detected phage RNA (**Figure 5A**). The intracellular abundance of the phage genes was comparable in the three *L. monocytogenes* strains under study (**Figure S6A**), indicating that Zea specifically affects the quantity of extracellular RNAs and not their expression level. However, when comparing *wt* and *Δzea* strains, we did not find remarkable changes in the amount of extracellular phage RNA. This is probably due to the low expression level of Zea by *wt* bacteria in MM, as revealed by qPCR (**Figure S5**). We next evaluated the abundance of another class of highly enriched RNAs specifically bound to Zea: the lma-monocin RNAs (**Figure S3**). The lma-monocin locus is considered to be a cryptic prophage specific of the *Listeria* genus whose function remains elusive (*20, 21*). Overexpression of Zea increased the amount of the lma-monocin RNAs in the culture medium (**Figure 5B**), mirroring the results obtained for the A118 phage. Also in this case, the intracellular level of lma-monocin RNA was comparable among the three *L. monocytogenes* strains used (**Figure S6B**).

**Figure 5.**
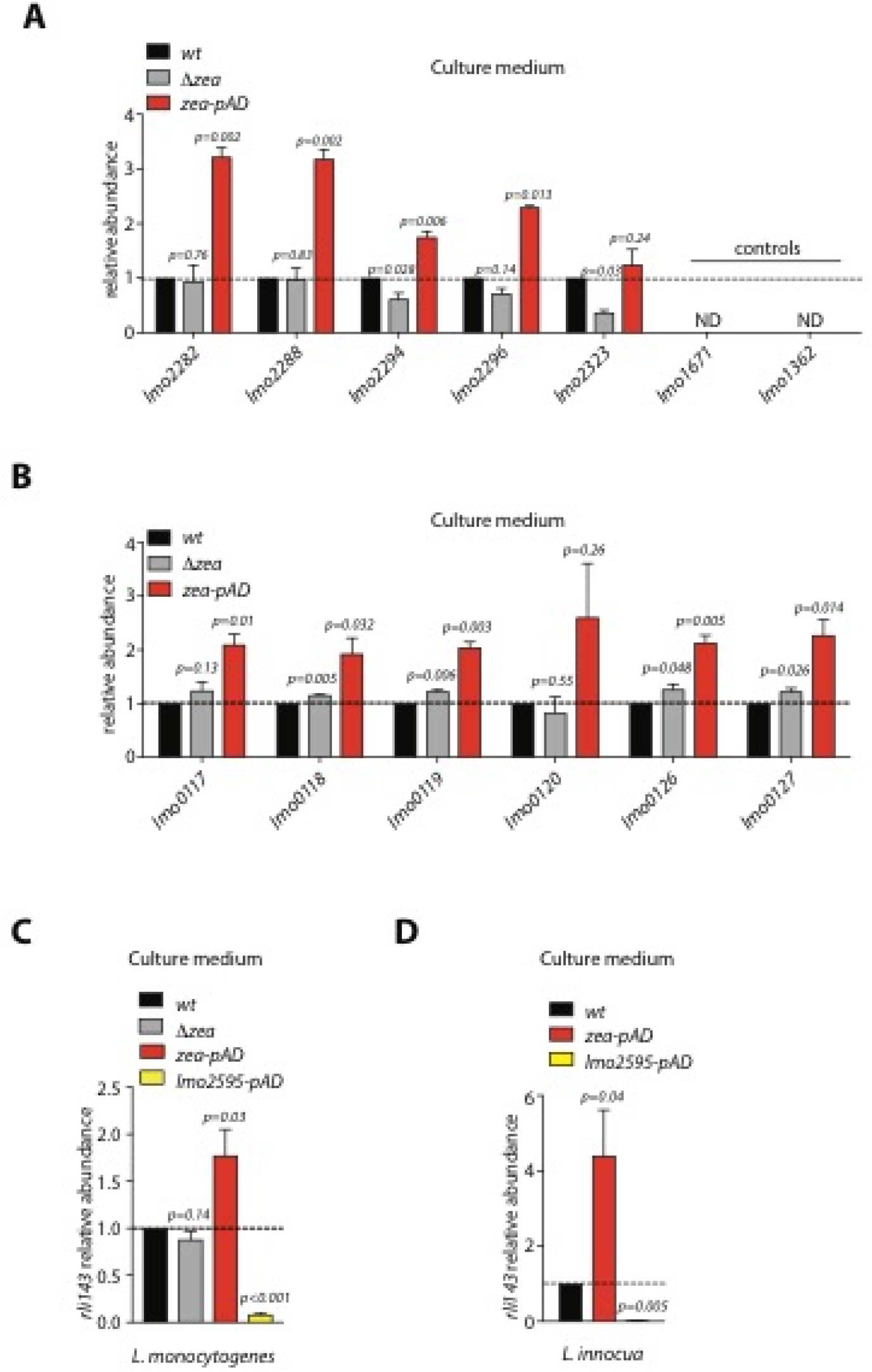
Zea controls the abundance of its target RNAs in the culture medium. qRT-PCR analysis on RNA extracted from the culture medium of different *L. monocytogenes* strains (as indicated) for: **(A)** selected phage and control genes; (B) the lma-monocin locus; **(C)** *rli143* in *L. monocytogenes*; **(D)** *rli143* in *L. innocua.* The relative abundance was calculated after normalization to the *wt* sample. The dashed lines indicate the relative abundance in the *wt* strain. Values reprents means ± SEM, n=3. ND, not detected. Statistical significance determined by two tailed *t*-test.

This approach could not be applied to secreted small RNAs detected by RIP-Seq due to their low expression levels. To overcome this problem, we overexpressed *rli143* in wt, *Δzea* and *zea-pAD L. monocytogenes* and then measured *rli143* abundance in the culture medium. As a control, we generated a fourth strain overexpressing both *rli143* and Lmo2595, another secreted protein of *L. monocytogenes* that, like Zea, harbors a canonical signal peptide (*10*). In line with the above results, *rli143* accumulated in the culture medium when Zea was co-overexpressed (**Figure 5C**) but not when Lmo2595 was co-overexpressed. The intracellular expression level of *rli143* was comparable in all of the strains (**Figure S6C**).

To further establish a role for Zea in the regulation of the extracellular amount of RNA, we generated a *L. innocua* strain overexpressing either *rli143* alone, or *rli143* with Zea, and then measured the abundance of *rli143* in the culture medium. As a control, *rli143* was co-expressed with Lmo2595. Of note, Zea and Lmo2595 are *L. monocytogenes* proteins that are also secreted when expressed in *L. innocua* (data not shown). We found that co-expression of Zea and *rli143* induced an even greater accumulation of *rli143* in the culture medium compared to *L. monocytogenes* (**Figure 5D**). The intra-bacterial abundance of *rli143* was comparable in all the *L. innocua* strains (**Figure S6D**). Altogether, these data show that the overexpression of Zea induces accumulation in the culture medium of phage-derived and small RNAs that are Zea-binding RNA species.

Zea overexpression could increase the amount of extracellular RNA by promoting its export from bacteria and/or by promoting its stabilization in the culture medium. Interestingly, we found that Zea protected *rli143* from RNase-mediated degradation *in vitro* (**Figure S7**) indicating that the mechanism of RNA stabilization may partially account for the increased amount of extracellular RNA.

As a last approach to definitively establish the impact of Zea on secreted RNA, we performed RNA-Seq analysis on extracellular RNAs prepared from the *wt* and *zea-pAD* strains. We reasoned that, if Zea increases the secretion and/or protection of a specific subset of extracellular *L. monocytogenes* RNAs, then its overexpression should increase the overall amount of those RNAs in the medium. We thus purified extracellular RNA from three independent samples (3 from *wt* and 3 from *zea-pAD)* and performed sequencing. Differential gene expression analysis revealed that, besides the overexpressed Zea RNA, 36 endogenous transcripts were significantly more abundant in the medium of the *zea-pAD* strain (**Figure S8A**). The vast majority of these transcripts were mRNAs (78%), while a small percentage represented sRNAs and antisense RNAs (10% and 8%, respectively) (**Figure S8A**). We then examined the correlation between Zea overexpression and the higher amount of extracellular RNA; we intersected the differential extracellular abundance dataset with the dataset of the Zea RIP-Seq experiment performed in the culture medium (i.e. the RNAs in complex with Zea). Strikingly, we found that one third (12 RNAs out of 37) of the transcripts enriched in the culture medium when Zea was overexpressed was also associated with Zea in the RIP-Seq dataset (**Figure S8B**). This indicates that a subset of transcripts found in complex with Zea becomes more abundant in the medium following Zea overexpression (exact right rank fisher’s test: *P* = 7.18×10^−5^). Of note, among these 12 enriched secreted RNAs, 8 RNAs proceeded from the A118 phage. RT-qPCR analysis of intrabacterial phage RNA from the *wt* and *zea-pAD* strains revealed similar amounts of the majority of the phage genes tested, indicating that Zea does not affect the expression of phage genes (**Figure S8C**). Altogether, our results show that Zea binds a subset of *L. monocytogenes* RNAs and that its overexpression increases their abundance in the extracellular medium.

## Zea modulates *L. monocytogenes* virulence

The absence of a Zea ortholog in *L. innocua* (**Figure 1A**), prompted us to assess whether Zea could impact *L. monocytogenes* virulence. Of note, the *Δzea* strain grew as well as the *wt* strain both in broth and in tissue cultured cells (macrophages and epithelial cells; data not shown). We thus examined the properties of the *wt* and *Δzea* strains in a mouse infection model. After intravenous inoculation, the *Δzea* strain showed a significant increase in bacterial load after 72h both in the liver (**Figure 6A**) and in the spleen (**Figure 6B**). These results indicate that Zea is an effector that dampens *L. monocytogenes* virulence.

**Figure 6.**
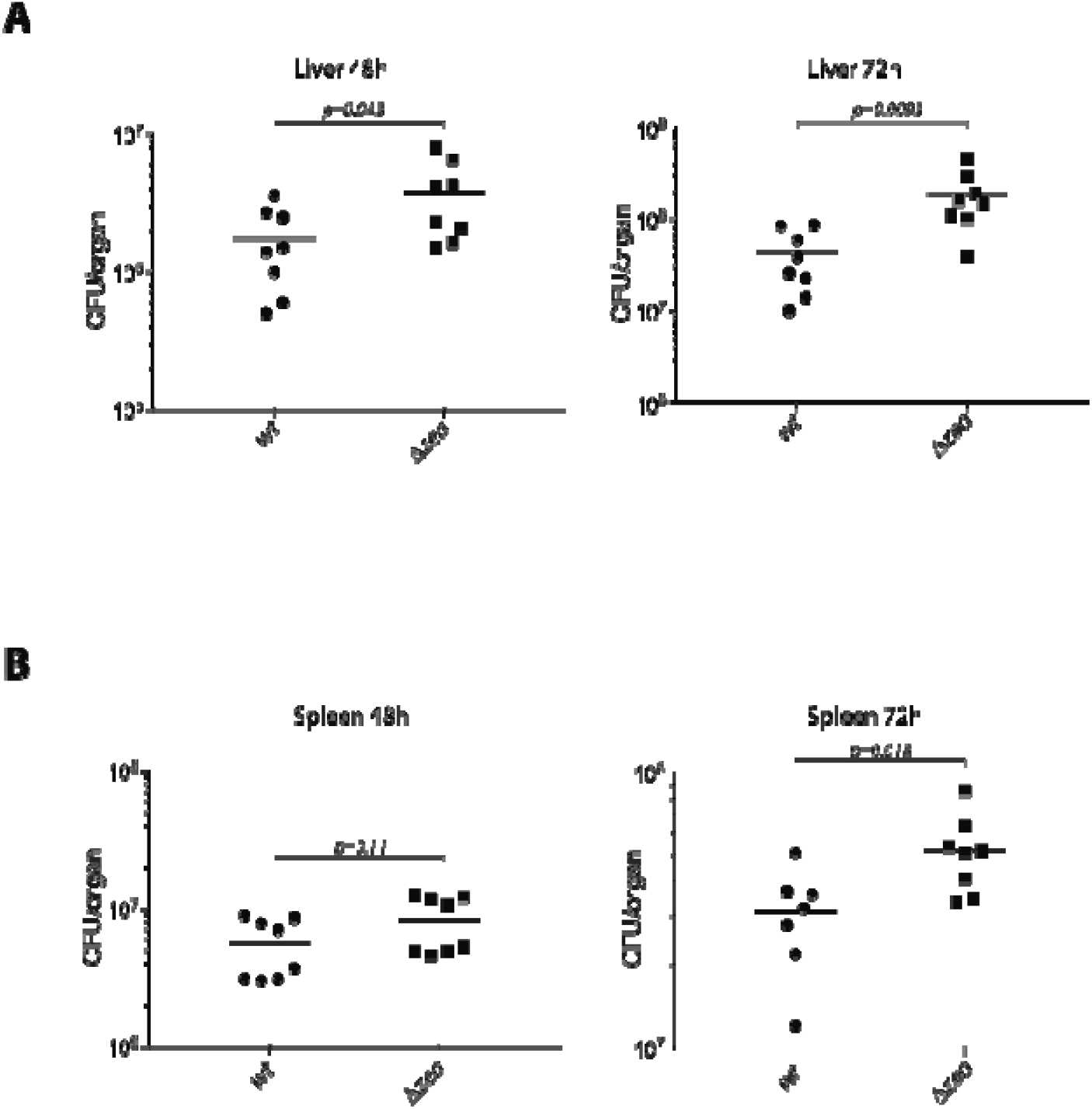
Zea regulates *L. monocytogenes* virulence. Balb/c mice were inoculated intravenously with 1 × 10^4^ CFUs of the *L. monocytogenes* EGD-e (wt) or the strain deleted of the *zea* gene *(Δzea).* After 48 h and 72 h post-infection, livers **(A)** and spleens **(B)** were recovered and CFUs per organ were assessed by serial dilution and plating. The number of bacteria in each organ is expressed as log10 CFUs. Black circles and squares depict individual animals. The lines denote the means ±SEM, n=2. Statistical significance determined by two tailed *t*-test.

## Zea potentiates the type I IFN response in a RIG-I dependent fashion

Three RBPs of the RIG-I-like receptor (RLR) family (RIG-I, MDA5 and LGP2) can sense nonself RNA in the cytoplasm but only RIG-I and MDA5 can trigger the type I IFN signaling cascade (*22*). A recent approach has successfully helped to identify viral RNA sequences bound either to RIG-I, MDA5 or LGP2 during viral infections (*23, 24*). This method is based on the affinity purification of stably expressed Strep-tagged RLRs followed by sequencing of their specific viral RNA partners. We thus applied this approach to obtain *L. monocytogenes-specific* RNAs bound to each of the RLRs upon infection with *L. monocytogenes wt.* We infected HEK293 cells stably expressing Strep-tagged RLRs (or Strep-tagged mCherry as a negative control) with *L. monocytogenes wt* and pulled down the Strep-tagged proteins. Co-purified RNA molecules from three independent replicates were sequenced and mapped to the *L. monocytogenes* genome. We found 15 RNAs specifically enriched in the RIG-I pull-down and 9 of them (60%) belonged to the phage A118 locus (**Figure S9**). We did not identify specific RNAs bound to MDA5 and LGP2, in agreement with a previously suggested major role of RIG-I, and a minor role of MDA5, in *L. monocytogenes-induced* IFN response infection (*25, 26*). These data indicate that, during infection, *L. monocytogenes* phage RNAs gain access to the host cytoplasm, where they specifically bind to RIG-I.

Since Zea is secreted and binds phage RNAs, we investigated whether it could participate in the RIG-I-dependent signaling. We first compared the expression of IFNβ in cells infected with *L. monocytogenes wt versus* cells infected with *zea-pAD*. qPCR analysis revealed that overexpression of Zea increased the amount of IFNβ while IFNγ was undetectable, as expected in non-immune cells (**Figure 7A**). Zea overexpression did not increase the expression of the proinflammatory cytokine interleukin 8 (IL8). Thus, Zea can specifically activate a type I IFNβ response. Next, to address whether the increased IFN response was mediated by RIG-I, we repeated the same experiment in RIG-I knock-down cells. The Zea-induced IFNβ stimulation was strongly impaired after RIG-I silencing (**Figure 7B**). These data indicate that Zea specifically enhances RIG-I-dependent type I IFN response.

**Figure 7.**
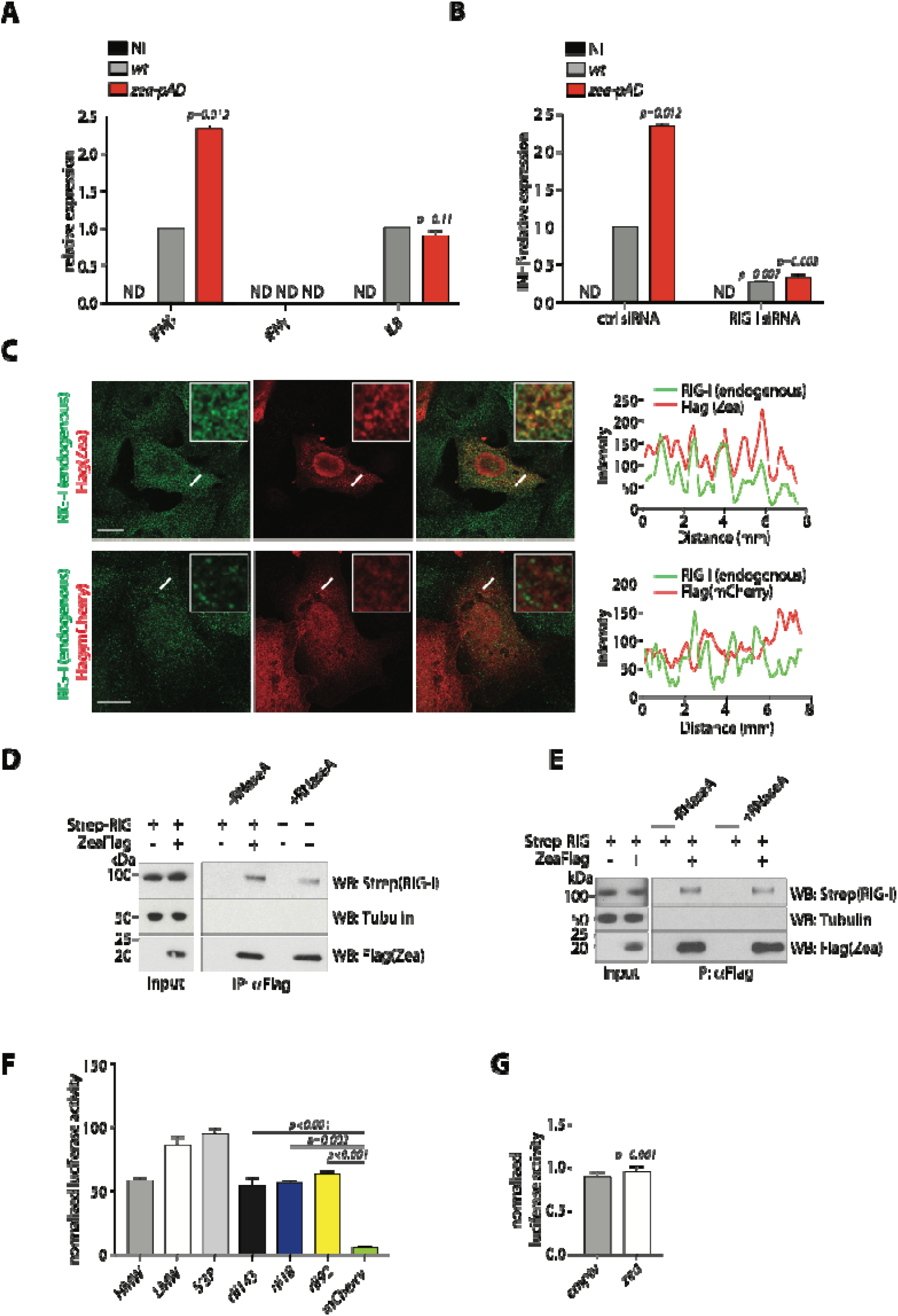
Zea interacts with RIG-I and stimulates a RIG-I dependent IFN response. **(A)** RT-qPCR analysis of *IFNβ, IFNγand* interleukin 8 *(IL8)* expression in response to infection with *wt* and *zea-pAD L. monocytogenes* in LoVo cells infected for 17 hours (MOI 5). The relative expression was calculated after normalization to (i) the GAPDH as a housekeeping gene and (ii) to the *wt* sample. ND, not detected. **(B)** RT-qPCR analysis of IFNβ expression in response to infection with *wt* and *zea-pAD L. monocytogenes* in LoVo cells transfected with control siRNA (ctrl siRNA) or with RIG-I targeting siRNA (RIG-I siRNA) and infected for 17 hours (MOI 5). Values reprents means ± SEM, n=3. Statistical significance was determined by two tailed *t*-test. **(C)** Representative confocal images of LoVo cells transfected with Flag-tagged Zea (top panel) or Flag-tagged mCherry (bottom panel), fixed and processed for immunofluorescence by using an anti-Flag antibody (red) and an anti-RIG-I antibody (green). The co-localization between Zea and RIG-I was assessed with a line scan (white line) whose fluorescence intensity is plotted in red for ZeaFlag and in green for RIG-I. Top right insets: magnification of the region in which the line scan was performed. Scale bars, 10 μm. **(D)** Representative Co-IP between Flag-tagged Zea and Strep-tagged RIG-I. Immunopurified ZeaFlag treated with RNase (+ RNaseA, 100 μg/mL) or untreated (-RNaseA) was incubated with a cell lysate from HEK-293 cells stably expressing Strep-tagged RIG-I (*23*). Input and immunoprecipitated materials (IP) were probed with antibodies against Flag-tag, Strep-tag and tubulin (which served as a negative control). **(E)** Representative Co-IP between Flag-tagged Zea and Strep-tagged RIG-I. LoVo cells were co-transfected with the plasmids encoding Flag-tagged Zea and Strep-tagged RIG-I and ZeaFlag was then immunoprecipitated and treated with RNase (+ RNaseA, 100 μg/mL) or untreated (-RNaseA) before elution with an anti-Flag peptide. Input and immunoprecipitated materials (IP) were probed with antibodies against Flag-tag, Strep-tag and tubulin (which served as a negative control). **(F)** The immunostimulatory activity of Zea-interacting small RNAs *(rli143, rli18* and *rli92)* was assessed by transfection into ISRE reporter cells lines (*29*). Values reprents means ± SEM, n=3. Firefly luciferase activity was measured and normalized to mock-transfected cells. *HMW* (high molecular weight), *LMW* (low molecular weight) and *5’3P* (5’ triphosphate-RNA) were used as positive controls. An *mCherry* RNA fragment served as a negative control. **(G)** The immunostimulatory activity of the Zea protein was assessed by transfection of a Zea-encoding plasmid (zea) into the ISRE reporter cells line (*29*). Transfection of an empty plasmid *(empty)* served as a negative control. Firefly luciferase activity was measured and normalized to mock transfected cells. Values reprents means ± SEM, n=3. Statistical significance was determined by two tailed ŕ-test.

The finding that Zea activates RIG-I signaling led us to examine whether the two proteins might share the same localization in cells. Attempts to detect endogenous Zea in infected cells by western blot or immunofluorescence were unsuccessful as our antibodies cross-reacted with unknown mammalian proteins. We thus infected cells with *zea^Flag^-pAD L. monocytogenes* and used an anti-Flag antibody for detection. Western blotting analysis of cytosolic and nuclear fractions prepared from infected cells revealed that Zea was present in both host cell compartments (**Figure S10A**) whereas RIG-I is mostly cytosolic (*27, 28*). Transfected Flag-tagged Zea also localized both to the cytoplasm and the nucleus. (**Figure S10B**). Thus, the fraction of Zea present in the cytosol might be compatible with the RIG-I-dependent signaling. We next tested whether Zea and RIG-I co-localized in cells. We found several loci of colocalization between transfected Flag-tagged Zea and endogenous RIG-I, indicating a spatial vicinity of the two proteins (**Figure 7C**). Of note, a negative control Flag-tagged mCherry protein did not co-localize with RIG-I (**Figure 7C**). These results prompted us to directly test whether Zea could interact with RIG-I. We immunopurified Zea from the culture medium of a *zea^Flag^-pAD* strain and used lysates of HEK293 cells stably expressing Strep-tagged RIG-I as a source of RIG-I protein (*23*). Immunopurified Zea efficiently and reproducibly pulled down Strep-tagged RIG-I from cell lysates, indicating specific interaction between the two proteins (**Figure 7D**). This interaction appeared partially stabilized by the presence of *L. monocytogenes* RNA, as pre-treatment of immunopurified Zea with RNaseA, reduced Zea/RIG-I binding, without abolishing it (**Figure 7D**). In agreement with a minor role of RNA in the Zea/RIG-I interaction, Flag-tagged Zea expressed after transfection in mammalian cells (which is therefore not bound to *L. monocytogenes* RNA) was able to interact with co-expressed Strep-tagged RIG-I, independently of RNA presence (**Figure 7E**). Altogether, our results show that Zea interacts with RIG-I and activates RIG-I-dependent type I IFN response.

Since RIG-I activation implies RNA binding, we sought to determine more precisely whether Zea-bound RNAs could trigger an IFN response. We used a reporter cell line stably transfected with a luciferase gene under the control of a promoter sequence containing five IFN-stimulated response elements (ISRE) (*29*). We found that transfection of the *in* vitro-transcribed Zea-interacting small RNAs showed strong immunostimulatory activity, while an mCherry control transcript did not (**Figure 7F**). This suggests that Zea can induce RIG-I activation in infected cells *via* its associated bacterial RNAs. The expression of Zea protein alone failed to induce any stimulation, indicating that, despite its capability to physically interact with RIG-I, Zea cannot promote RIG-I activation by itself (**Figure 7G**). We conclude that during infection, Zea interacts with and activates RIG-I and this activation likely depends on Zea-bound RNA (**Figure 8**).

**Figure 8.**
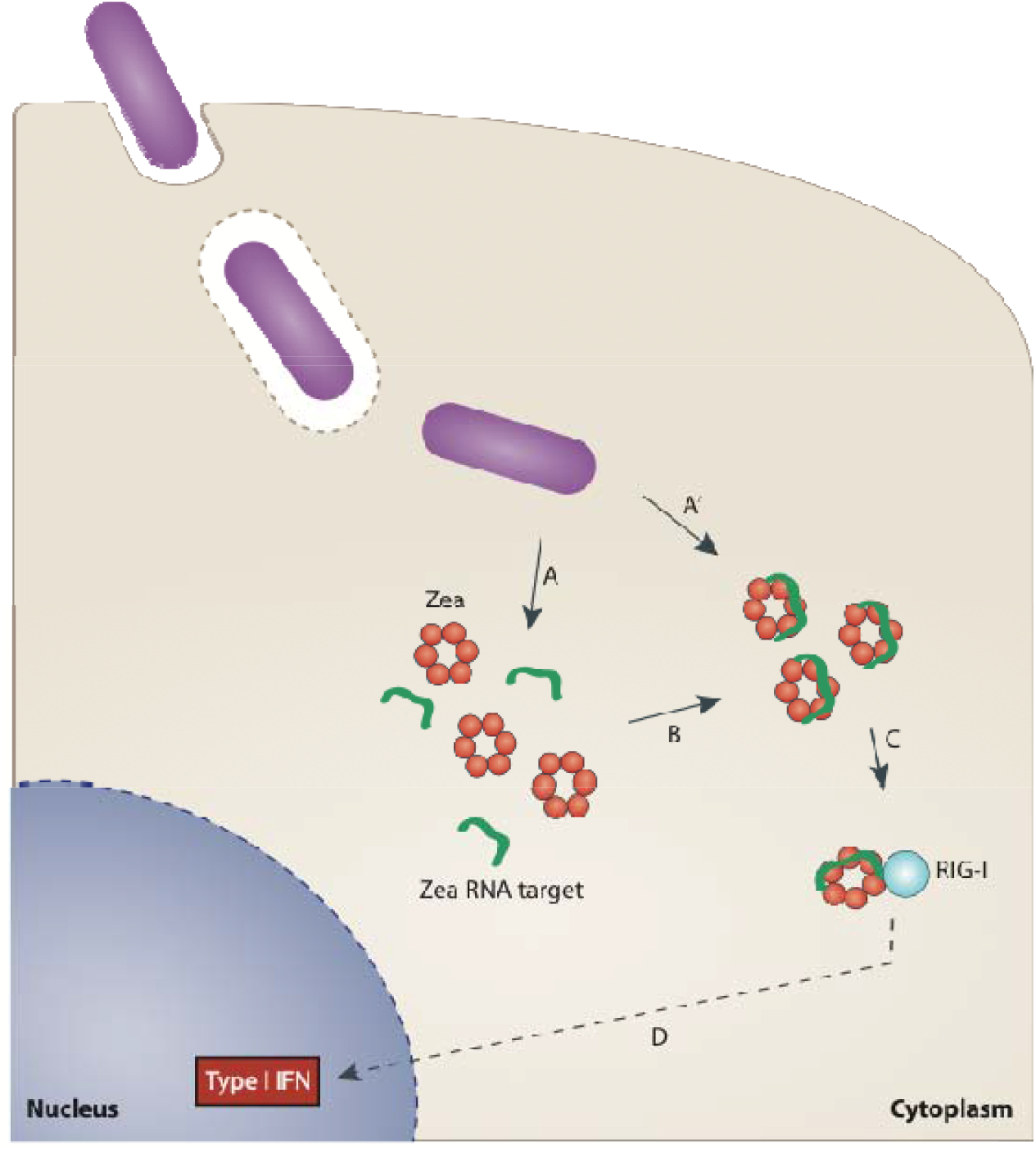
Proposed model for Zea-mediated regulation of IFN response. During infection, *L. monocytogenes* secretes Zea and RNA into the cytoplasm (A) which then assemble to form a ribonucleoprotein complex (B); in alternative, Zea can be directly secreted in an RNA-bound form (A’). Zea-containing ribonucleoprotein complexes then bind RIG-I (C), triggering a signaling cascade which would then activate the type I IFN response (D).

## Discussion

A key major finding of this study is the identification of the first secreted RBP from bacteria. We show that *L. monocytogenes* secretes Zea, a small RBP, which associates extracellularly with a subset of *L. monocytogenes* RNAs enriched in the phage and phage-derived RNAs. We found that the overexpression of Zea correlates with an increased amount of its RNA ligands in the culture medium. This might be due to two, non-mutually exclusive, phenomena, (i) increased secretion of Zea-RNA complexes, (ii) increased stabilization of Zea RNA targets following Zea accumulation in the extracellular medium. In the latter case, the pathway regulating RNA secretion would remain to be identified. The presence of Zea orthologues in other bacterial species indicates that the secretion of RBPs is a conserved phenomenon in prokaryotes. Our results suggest that additional RBPs remain to be identified in bacteria; however, their identification might be difficult as bacterial RBPs are likely devoid of classical RBDs (*8*). In agreement with this hypothesis, the analysis of the Zea protein sequence showed that Zea does not possess any recognizable RNA-binding region.

Our work reveals a novel mechanism regulating the induction of the INF response during bacterial infection which involves the secretion of an RBP. We provide evidence that during *L. monocytogenes* infection, a Zea-containing ribonucleoprotein complex binds to RIG-I and stimulates RIG-I activation. In agreement with this hypothesis, Zea devoid of its RNA targets cannot activate RIG-I, while Zea bound RNAs are able to do so. In addition, Zea and RIG-I bind a similar subset of *L. monocytogenes* RNAs, which is enriched in phage RNAs. This study provides the first identification of the bacterial RNA species recognized by RIG-I during bacterial infection and uncovers that RIG-I binds viral RNAs from an invading bacterium. These data reinforce the concept that RIG-I has primarily evolved to sense viral RNAs and highlight that phage RNAs from a bacterial pathogen contribute to RIG-I activation.

Our data show that Zea dampens *L. monocytogenes* virulence *in vivo* as its deletion results in an increased bacteria burden in the organs of infected mice. Strikingly and in agreement with our results, Zea is absent in the hypervirulent strains of *L. monocytogenes* (lineage I), while it is well conserved in the strains from lineage II, which includes a smaller number of clinical isolates. This indicates that Zea is a factor that, when present, contributes to render *L. monocytogenes* less virulent. Given the complexity of the innate immune response to *L. monocytogenes* infection (*30–34*), it is difficult to establish the precise role of Zea *in vivo.* We show that Zea participates in the modulation of the IFN response, but we do not exclude that Zea might have additional roles during infection. We found that Zea also localizes to the nucleus of infected cells (**Figure 7C and Figure S10**), possibly to affect host nuclear functions. It is conceivable that the phenotype observed *in vivo* is a consequence of these additional features of Zea. Given the striking ability of Zea to bind RNA, one important question to address in the future is whether Zea could also bind mammalian RNA during *L. monocytogenes* infection. Notably, Zea orthologs are also present in other bacteria, that normally reside in the environment and are rarely associated with disease. We do not exclude that, in addition to its role in *L. monocytogenes* virulence, Zea might also play a role during the saprophytic life of *L. monocytogenes*. Of note, most of the bacterial species harboring *zea* orthologs are genetically tractable, making the elucidation of the role of Zea-interacting RNAs in environmental bacteria to be within reach. In conclusion, this study revealed the existence of an extracellular ribonucleoprotein complex from bacteria and highlights a novel mechanism involved in the complex host cell response to infection.

## Acknowledgments

We thank all members of the Cossart laboratory for helpful discussions. We thank Dr Birgitte Kallipolitis for providing the anti-Hfq antibody. We thank Edith Gouin for production of antibodies. We are grateful to R.Y. Sanchez David and R. Viber for technical support. We acknowledge the platforms of the Grenoble Instruct center (ISBG; UMS 3518 CNRS-CEA-UGA-EMBL) with support from FRISBI (ANR-10-INSB-05-02) and GRAL (ANR-10-LABX-49-01) within the Grenoble Partnership for Structural Biology (PSB). We thank Jost Enninga, Javier Pizarro-Cerda, Mélanie Hamon, Olivier Dussurget, Hélène Bierne and Filipe Carvalho for critical reading of the manuscript and useful advices.

---

**Figure S1.**
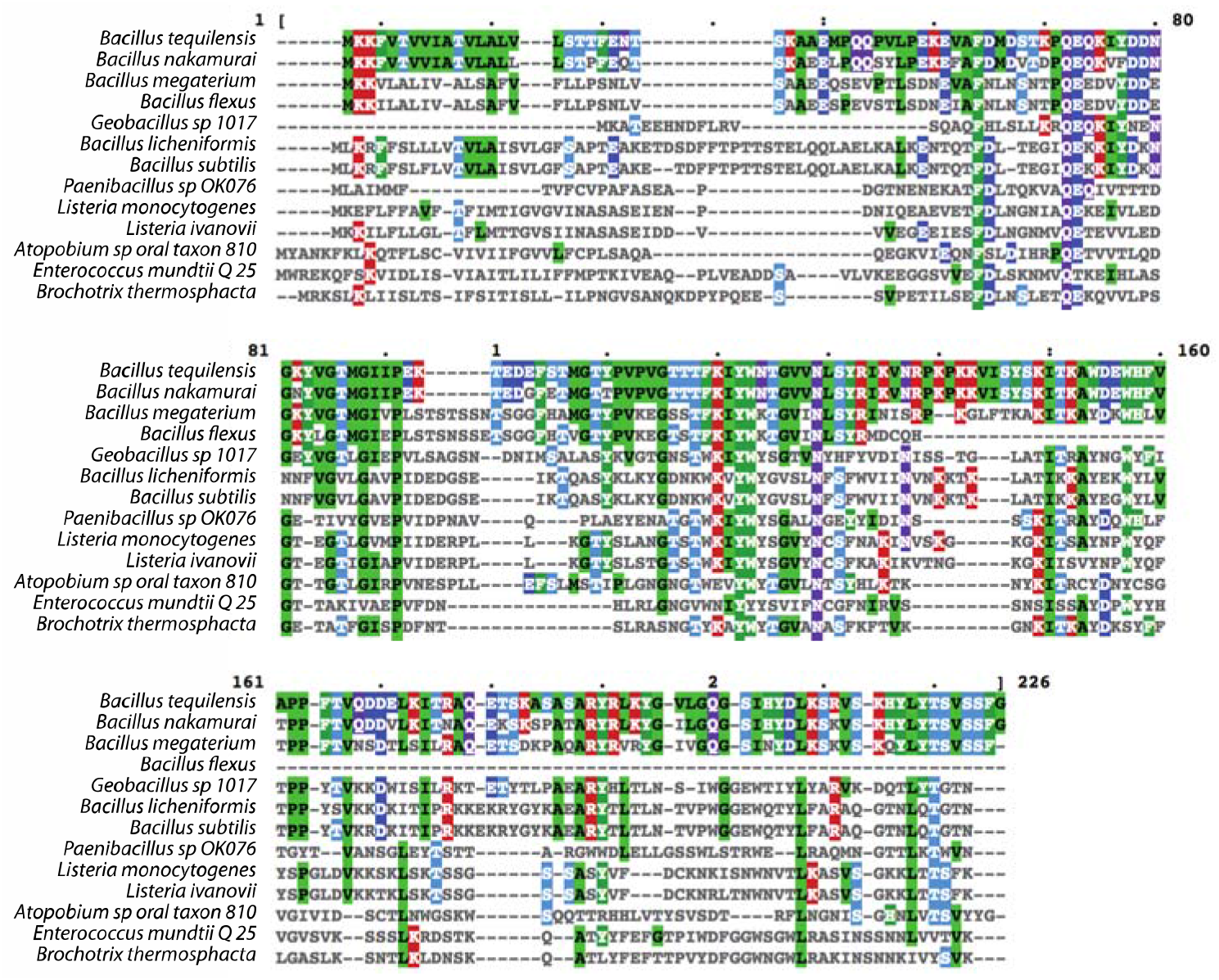
Protein sequence alignment of Zea orthologues. Multiple alignment of Zea orthologues performed with Clustal Omega. Zea-harboring species are indicated on the left. Conserved residues are highlighted with colors.

**Figure S2.**
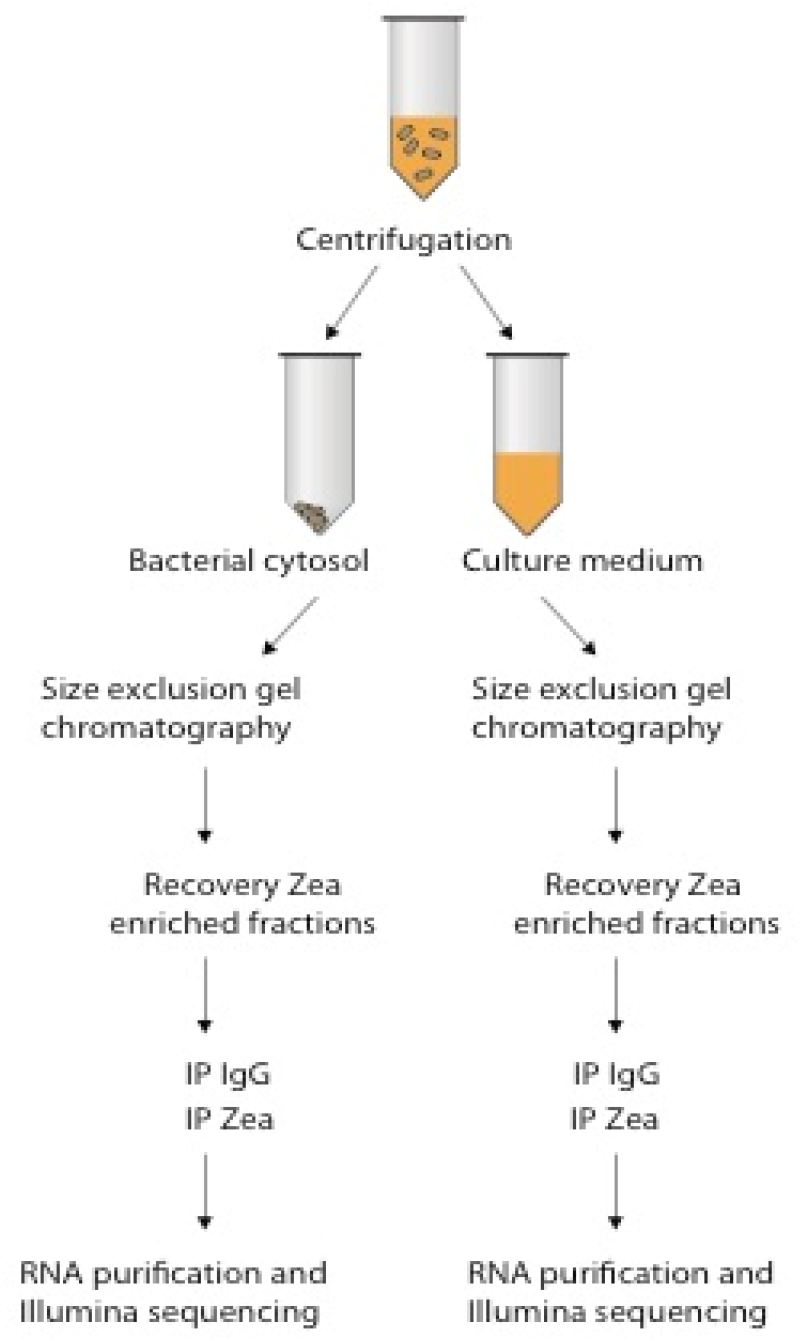
RIP-Seq protocol. Scheme representing the RIP-Seq protocol employed to identify the Zea-interacting RNAs (see Methods).

**Figure S3.**
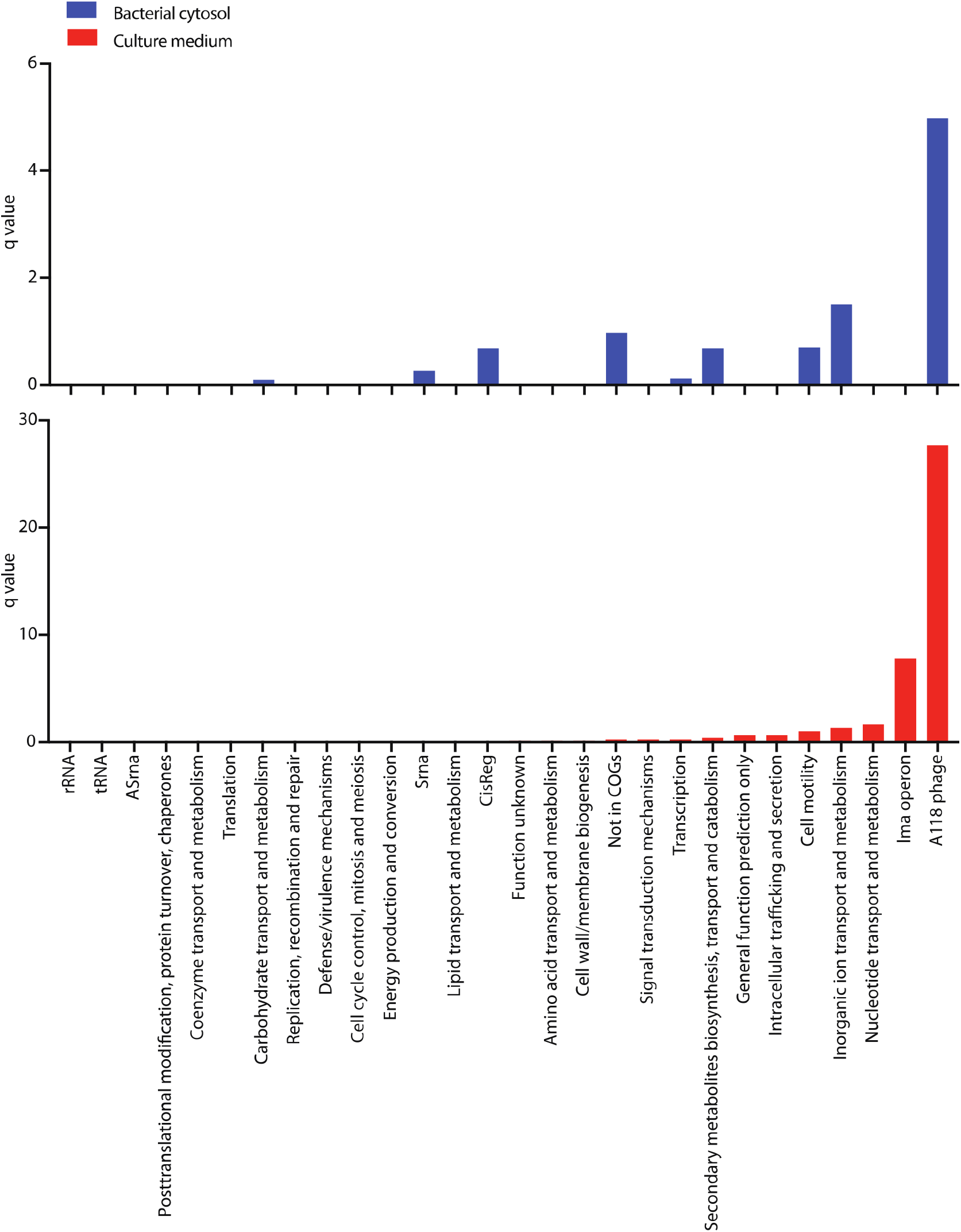
COG analysis of Zea-interacting RNAs. COG functional classes were assigned to RNAs identified *via* RIP-Seq in the bacterial cytosol and culture medium, as indicated. Fisher’s exact right rank test was run and q-values [-log_10_(p-value)] were calculated to detect overrepresented COG categories.

**Figure S4.**
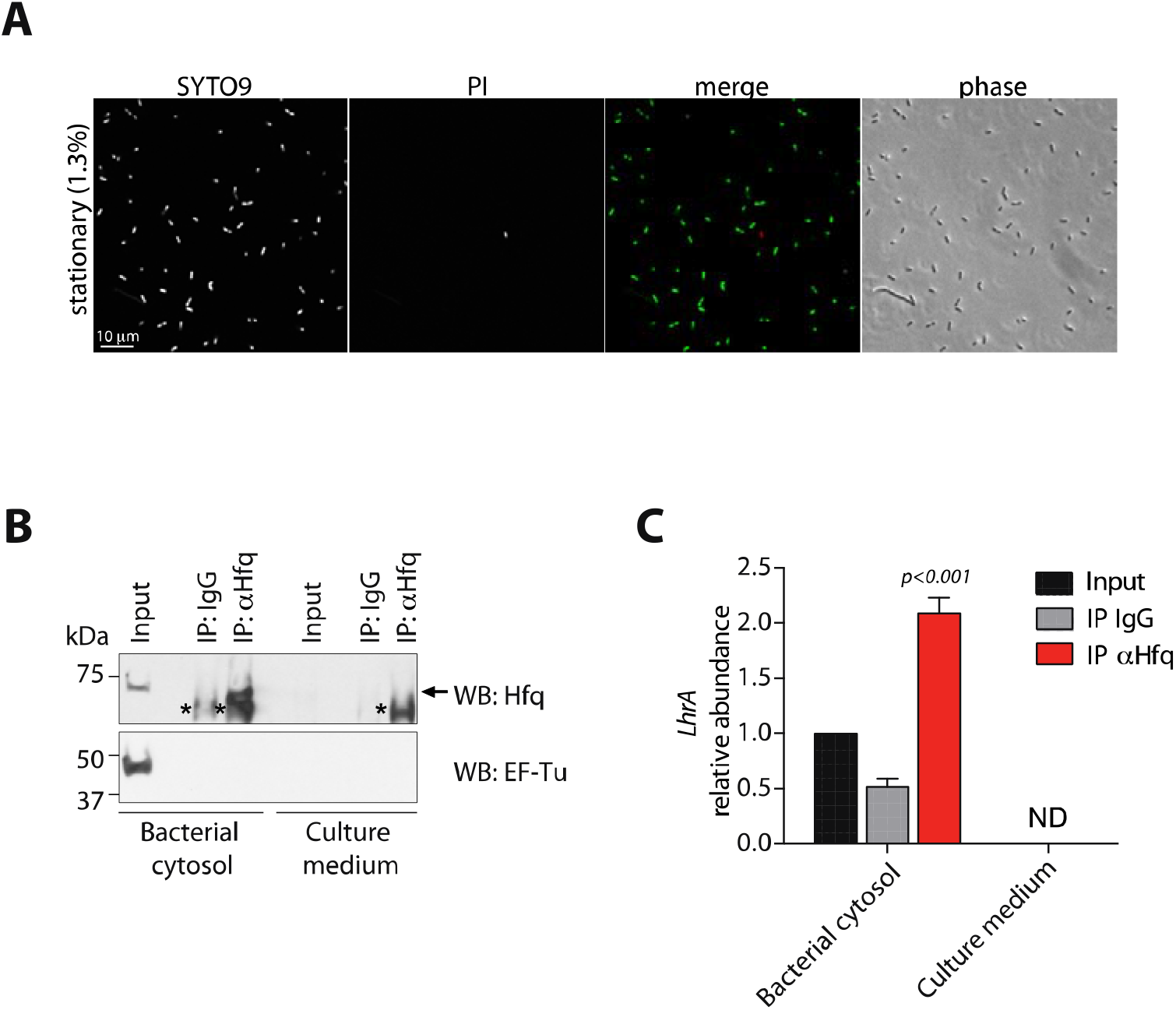
Extracellular Zea-bound RNAs do not derive from bacterial lysis. (A) Representative confocal microscopy image of *L. monocytogenes* viability by using live/dead staining assay. Live bacteria with intact cell membrane are stained with SYTO9 (green in the merged channel) and dead bacteria with damaged cell membrane are stained with propidium iodide (PI, red in the merged channel). The percentage of dead bacteria of one representative experiment is indicated on the left of the panel. (B) Representative immunoprecipitation (IP) with an anti-Hfq antibody of *L. monocytogenes* bacterial cytosolic extract and culture medium. WB with an anti-Hfq and anti-EF-Tu antibodies: starting material (Input), Hfq immunoprecipiated protein (IP:αHfq) and control immunoprecipitation (IP:IgG). Hfq runs as an hexamer (60kD). Asterisks and the arrow show the positions of the IgG heavy chain and Hfq, respectively. (C) RIP-qPCR for *LhrA* on RNA isolated from Hfq and control (IgG) immunoprecipitations in the bacterial cytosol and culture medium. The enrichment of *LhrA* was calculated after normalization to the input fraction. Values reprents means ± SEM, n=3. ND, not detected; statistical significance determined by two tailed t-test between the IgG IP and Hfq IP.

**Figure S5.**
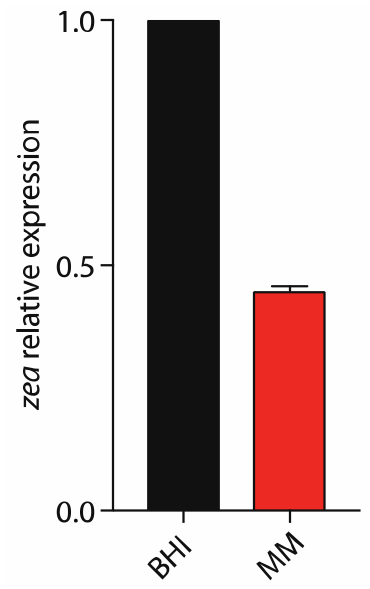
Zea is expressed at low level in minimal medium. qRT-PCR analysis on total RNA extracted from *L. monocytogenes wt* grown to the exponential phase in rich medium (BHI) or minimal medium (MM), showing low expression of Zea in MM. Data were normalized to *gyrA* mRNA and level in BHI. Values reprents means ± SEM, n=2.

**Figure S6.**
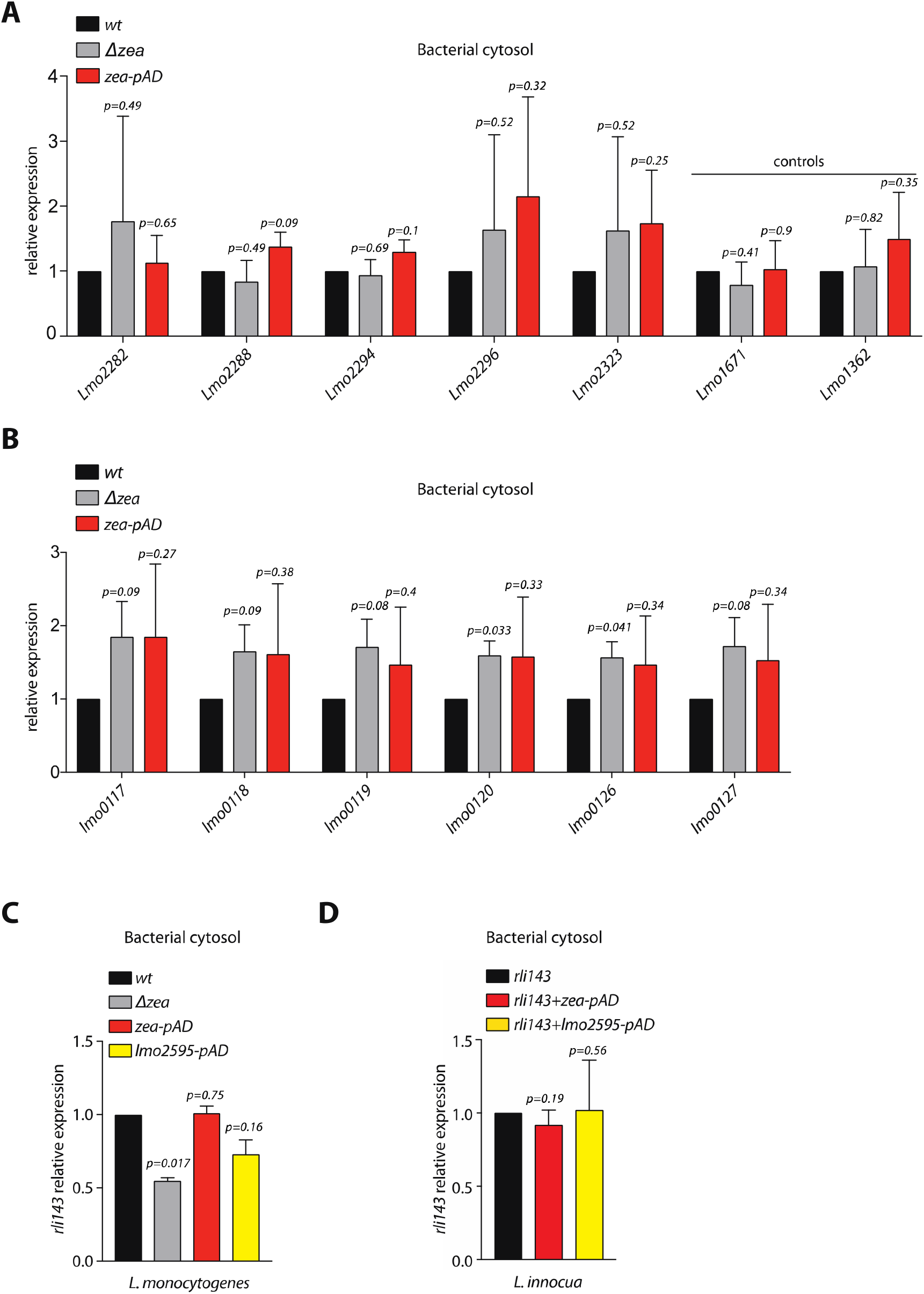
Phage A118, lma-operon and rli143 expression in *L. monocytogenes*. qRT-PCR analysis on total RNA extracted from the indicated *L. monocytogenes* strains grown to the exponential phase in MM. **(A)** Quantification of selected phage and control genes; **(B)** selected genes from the lma-monocin locus; **(C)** *r1i143* in *L. monocytogenes* and **(D)** *r1i143* in *L. innocua.* Data were normalized to *16S*rRNA and level in the *wt* strain. Values represents means ± SEM, n=3. Statistical significance determined by two tailed t-test.

**Figure S7.**
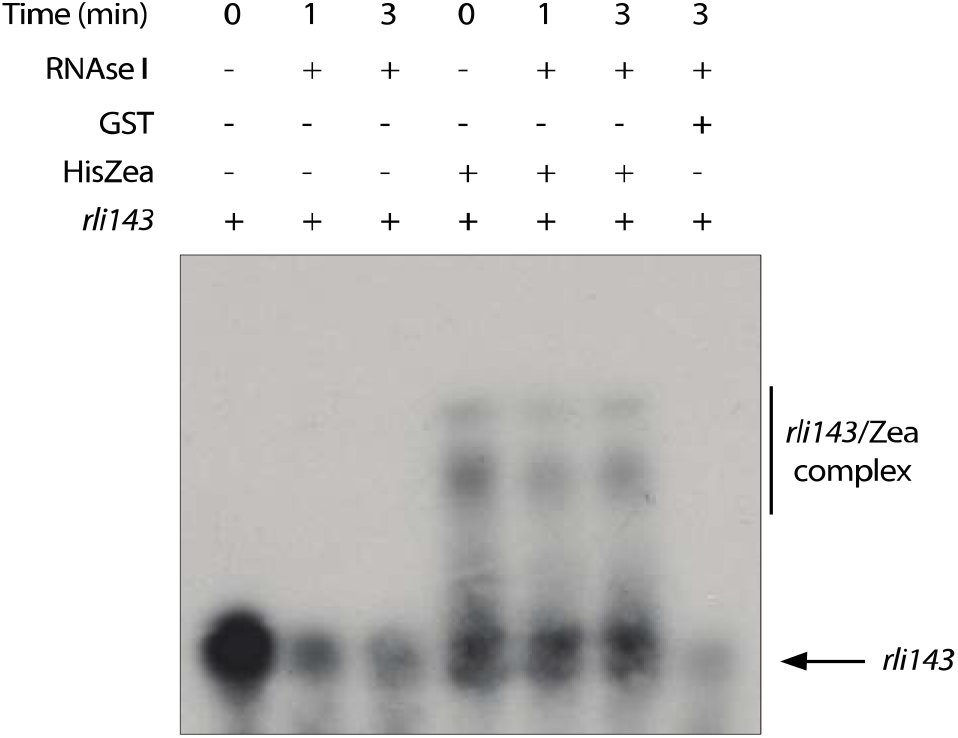
Zea protects rli143 from RNase degradation. *In* vitro-transcribed radiolabeled *rli143* was incubated with equimolar concentrations (1 μM) of either HisZea or GST (as a control). RNase I (0,0033U) was subsequently added and samples were incubated at 37 °C for the indicated time.

**Figure S8.**
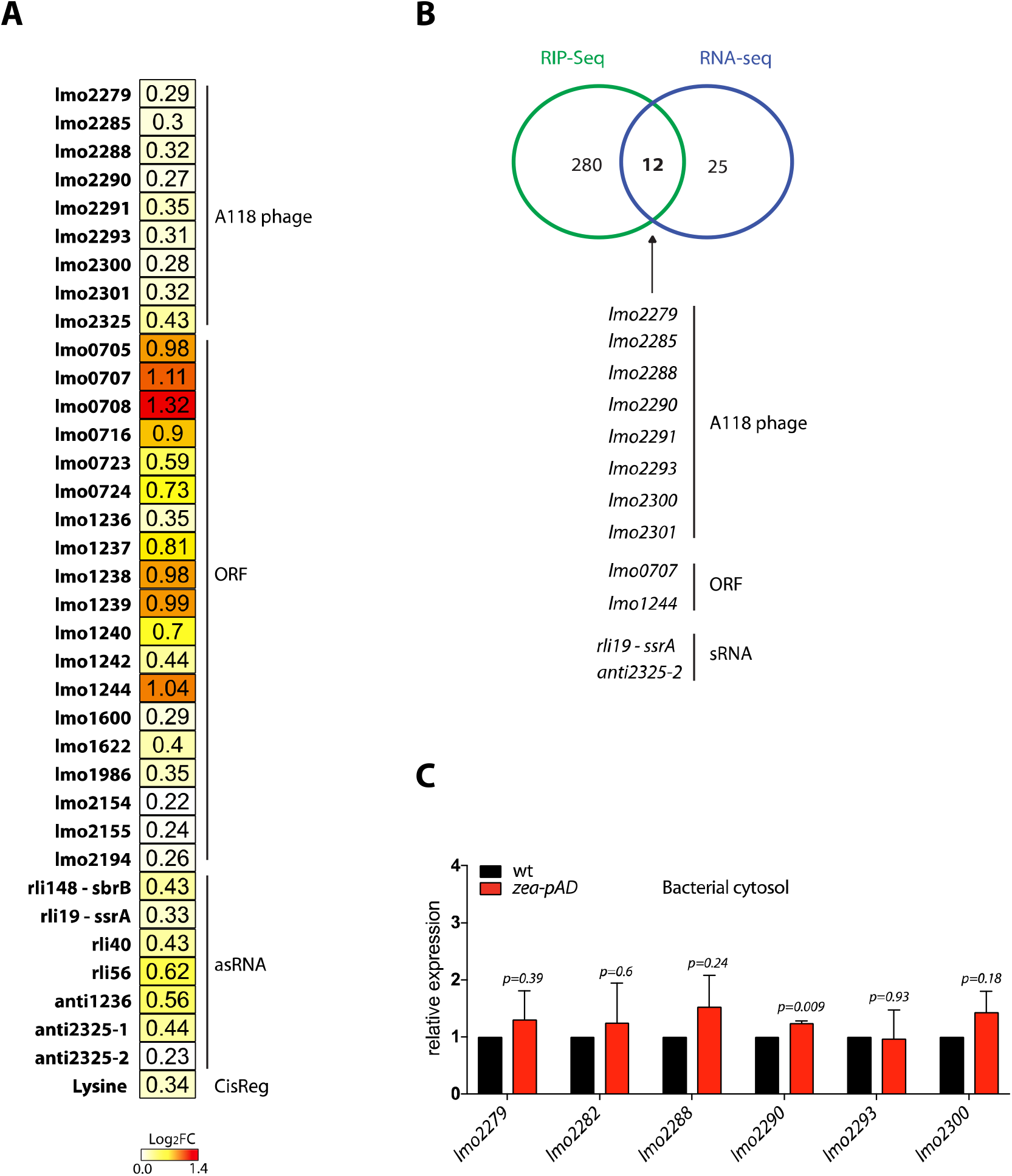
Zea regulates the extracellular amount of a subset of *L. monocytogenes* RNA. **(A)** Differential analysis of RNA-Seq quantification of secreted RNAs from *L. monocytogenes wt* and *Δzea+zea-pAD* strains (a zea-deleted strain in which one copy of the *zea* gene was integrated in the *L. monocytogenes* chromosome under the control of a constitutive promoter). The heat map shows 36 RNAs that are significantly enriched in the culture medium of *Δzea+zea-pAD* strain. Values in the squares are expressed as log_2_FC. **(B)** Overlap between the RIP-Seq and the RNA-Seq datasets. 12 RNAs are enriched in both the RIP-Seq and RNA-Seq, 280 are enriched only in the RIP-Seq, and 25 are differentially expressed only in the RNA-Seq, on a total of 3160 expressed L. *monocytogens* RNAs. A Fisher’s exact right rank test shows that the ratio of proportion of RNAs differently expressed and enriched in the RIP-seq is greater than 1 (p = 7.18×10^−5^). The list highlights the genes found in both datasets. **(C)** qRT-PCR analysis on total intracellular RNA extracted from *L. monocytogenes wt* and *Δzea+zea pAD* for the indicated phage genes. Data were normalized to *16S* rRNA. Values reprents means ± SEM, n=3. Statistical significance determined by two tailed t-test.

**Figure S9.**
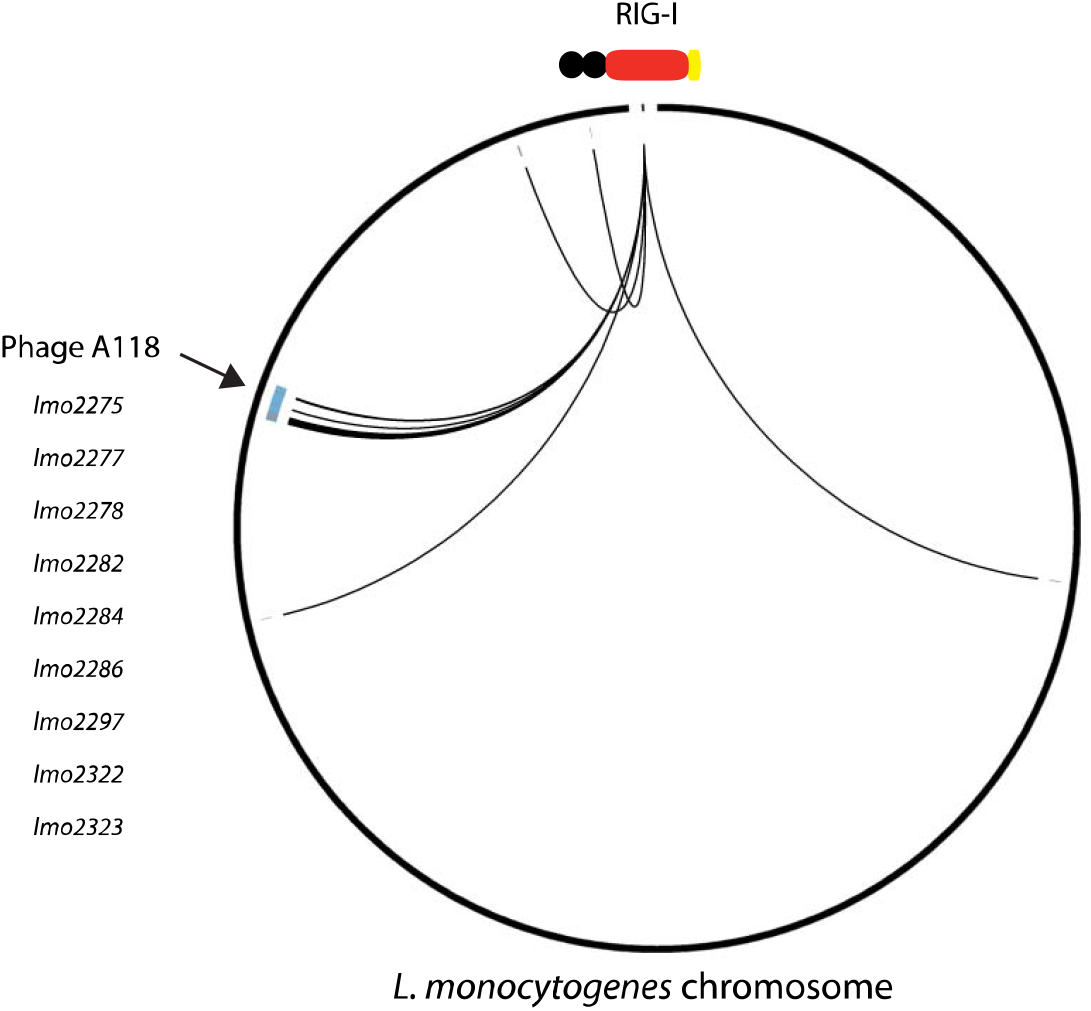
RIG-I binds the *L. monocytogenes* A118 phage RNA during infection. Interaction circo plot of *L. monocytogenes* RNAs specifically bound to RIG-I during *L. monocytogenes* infection. The circle represents the *L. monocytogenes* genome. A line from the schematized RIG-I to a genome locus indicates a preferential binding to a transcript compared to the negative control mCherry.

**Figure S10.**
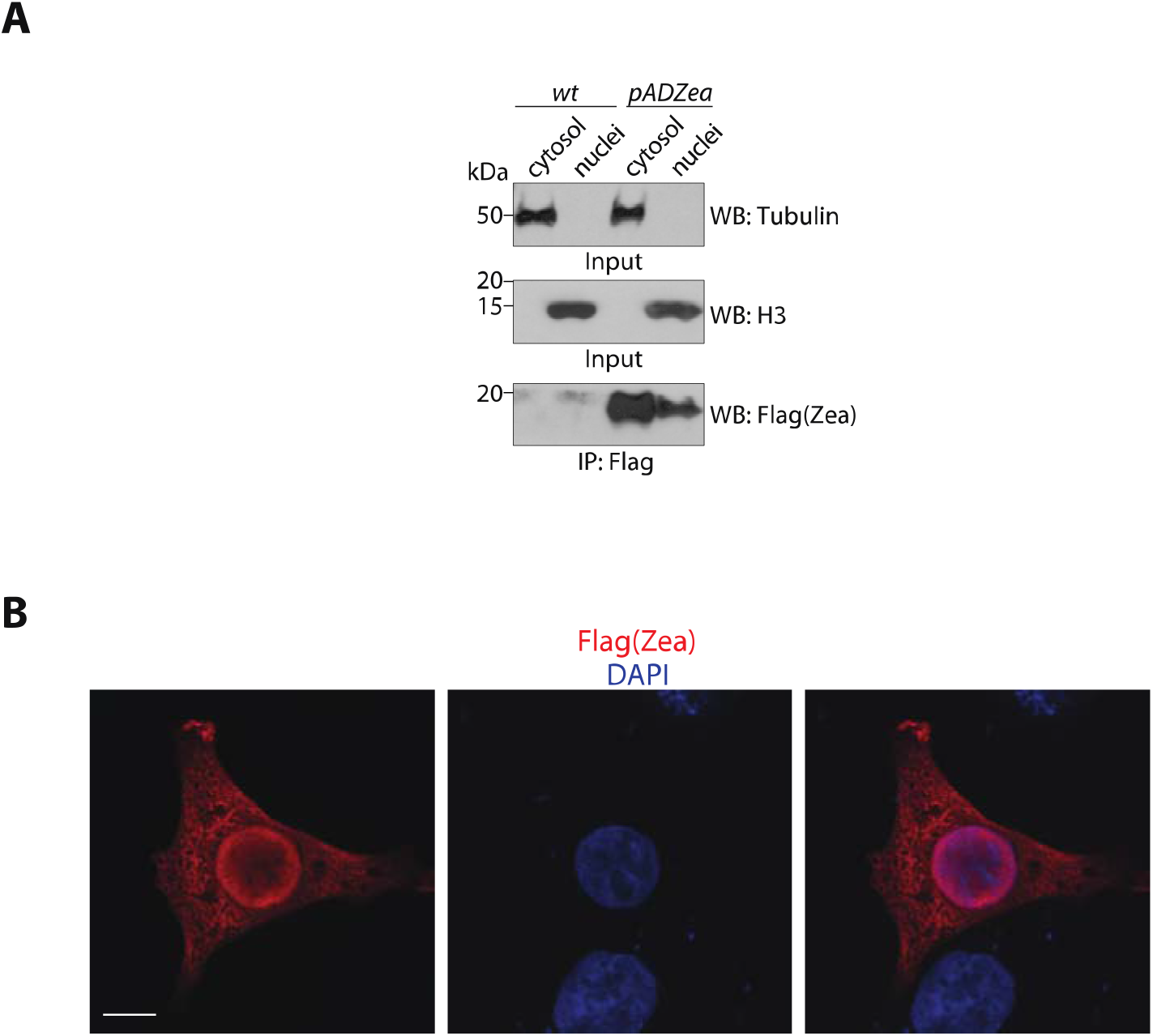
Zea localizes both to the host cell cytoplasm and nucleus. (A) IP analysis with an anti-Flag antibody of cytosolic and nuclear fractions from LoVo cells infected for 6 h with *wt* and *zea^Flag^-pAD* bacteria. Representative WB (antibodies as indicated) of total cytosolic and nuclear fractions (Input) and immunoprecipitated proteins (IP). (B) Representative confocal images of LoVo cells transfected with Flag-tagged Zea, fixed and processed for immunofluorescence by using an anti-Flag antibody (red); nuclei are stained with DAPI. Scale bar, 10 μm.

**Table S1.** Identification of Zea binding RNAs. The table shows the percentage of the different RNA species.

| BACTERIAL CYTOSOL | CULTURE MEDIUM |
| --- | --- |
| ORF: 89% (Phage 27.7%) | ORF: 94.5% (Phage 14.4%; Lma-monocin locus 3%) |
| ncRNAs: 11% | ncRNA: 5.5% |

**Table S2.** List of bacterial strains used in this study

|  |
| --- |
| *L. monocytogenes* EGD-e<br>(wt strain) |
| *L. monocytogenes* Δzea<br>(zea deleted strain) |
| *L. monocytogenes* Δzea+zea-pAD<br>(zea overexpressing strain in Δzea background) |
| *L. monocytogenes* zea-pAD<br>(zea overexpressing strain in wt background) |
| *L. monocytogenes* zea<sup>Flag</sup>-pAD+Zea<sup>HA</sup>-pP1<br>(zea<sup>Flag</sup> and Zea<sup>HA</sup> overexpressing strain in wt background) |
| *L. monocytogenes* zea<sup>Flag</sup>-pAD<br>(zea<sup>Flag</sup> overexpressing strain in wt background) |
| *L. monocytogenes* zea<sup>HA</sup>-pP1<br>(zea<sup>HA</sup> overexpressing strain in wt background) |
| *L. monocytogenes* rli143-pP1<br>(rli143 overexpressing strain in wt background) |
| *L. monocytogenes* rli143-pP1+zea-pAD<br>(rli143 and zea overexpressing strain in wt background) |
| *L. monocytogenes* rli143-pP1+lmo2595-pAD<br>(rli143 and lmo2595 overexpressing strain in wt background) |
| *L. innocua* rli143-pP1<br>(rli143 overexpressing strain in wt background) |
| *L. innocua* rli143-pP1+zea-pAD<br>(rli143 and zea overexpressing strain in wt background) |
| *L. innocua* rli143-pP1+lmo595-pAD<br>(rli143 and lmo2595 overexpressing strain in wt background) |

**Table S3.**
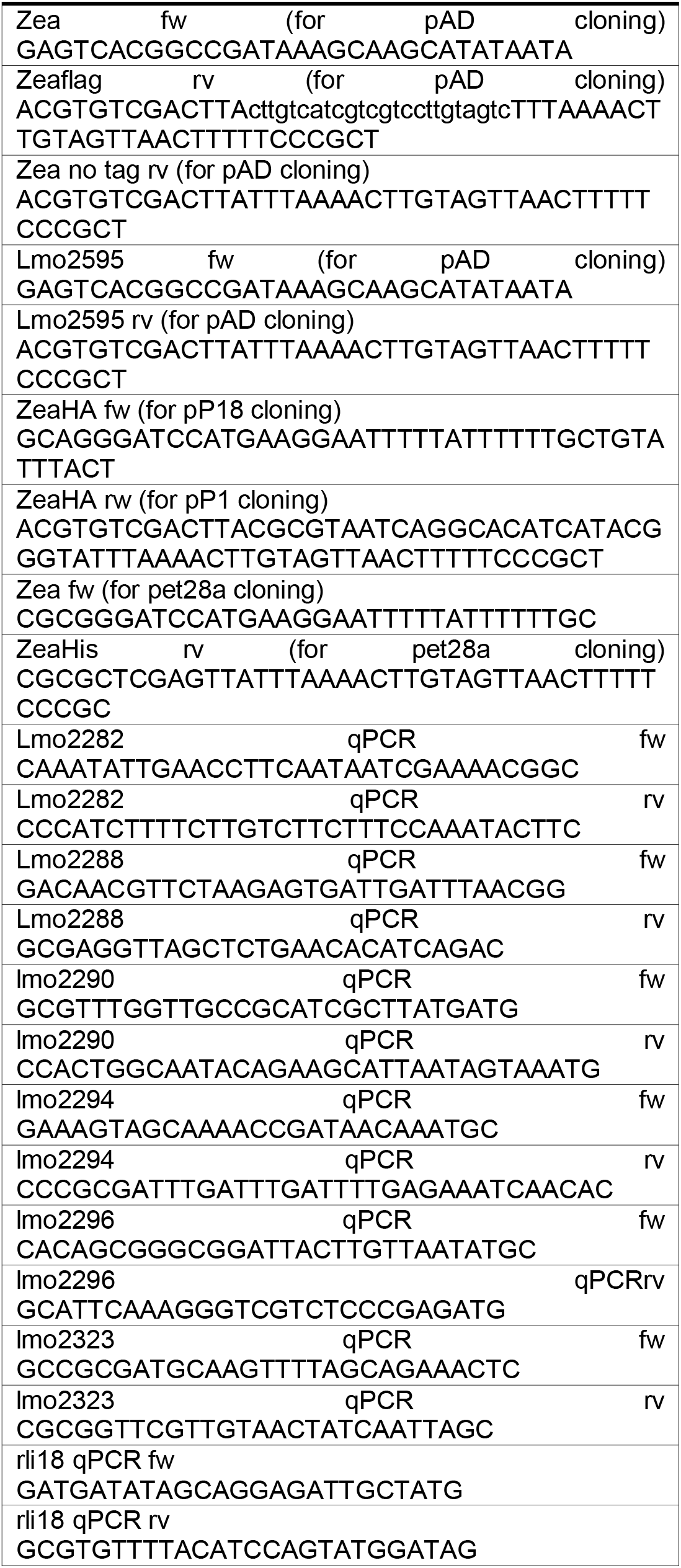
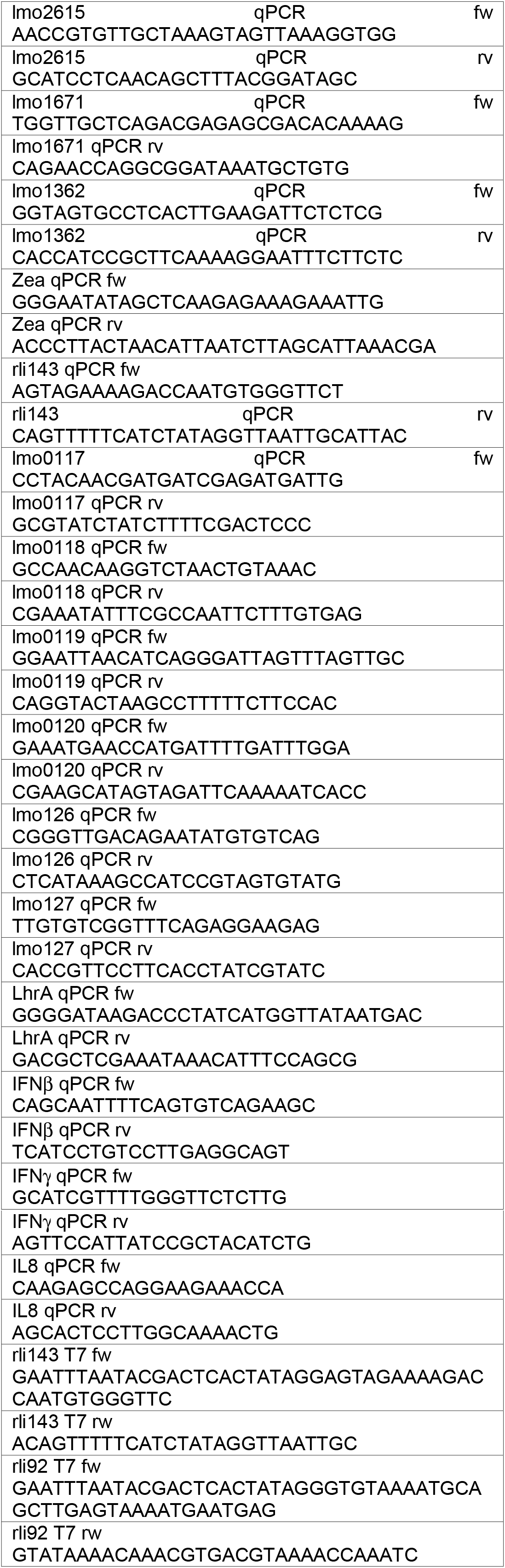
List of oligonucleotides used in this study

